# Secure Discovery of Genetic Relatives across Large-Scale and Distributed Genomic Datasets

**DOI:** 10.1101/2024.02.16.580613

**Authors:** Matthew M. Hong, David Froelicher, Ricky Magner, Victoria Popic, Bonnie Berger, Hyunghoon Cho

**Affiliations:** MIT, Cambridge, 02139, USA; Broad Institute of MIT and Harvard, Cambridge, 02142, USA; Yale University, New Haven, 06510, USA

## Abstract

Finding relatives within a study cohort is a necessary step in many genomic studies. However, when the cohort is distributed across multiple entities subject to data-sharing restrictions, performing this step often becomes infeasible. Developing a privacy-preserving solution for this task is challenging due to the significant burden of estimating kinship between all pairs of individuals across datasets. We introduce SF-Relate, a practical and secure federated algorithm for identifying genetic relatives across data silos. SF-Relate vastly reduces the number of individual pairs to compare while maintaining accurate detection through a novel locality-sensitive hashing approach. We assign individuals who are likely to be related together into buckets and then test relationships only between individuals in matching buckets across parties. To this end, we construct an effective hash function that captures identity-by-descent (IBD) segments in genetic sequences, which, along with a new bucketing strategy, enable accurate and practical private relative detection. To guarantee privacy, we introduce an efficient algorithm based on multiparty homomorphic encryption (MHE) to allow data holders to cooperatively compute the relatedness coefficients between individuals, and to further classify their degrees of relatedness, all without sharing any private data. We demonstrate the accuracy and practical runtimes of SF-Relate on the UK Biobank and All of Us datasets. On a dataset of 200K individuals split between two parties, SF-Relate detects 94.9% of third-degree relatives, and 99.9% of second-degree or closer relatives, within 15 hours of runtime. Our work enables secure identification of relatives across large-scale genomic datasets.

## Introduction

Collaborative studies that aim to jointly analyze genomic data from multiple parties are essential for increasing the sample sizes to enhance the discovery of biomedical insights. However, when sharing individual-level genetic data is not feasible due to privacy concerns (e.g., (Erlich et al. 2018)), the range of joint analyses that can be performed is severely limited. As a result, many existing collaborations have relied on simplified analysis pipelines where some key analysis steps, such as cohort identification, quality control procedures, and correction for confounding factors (e.g., population structure) are performed independently by each party on their respective datasets without considering the pooled data. This presents a key barrier to realizing the full potential of collaborative genomics research.

An important analysis task that is commonly omitted in collaborative studies is the identification of genetic relatives across isolated datasets. Identifying and excluding close relatives within a study cohort is a standard step in many genetic analyses (e.g., genome-wide association studies (Anderson et al. 2010)), because the presence of relatives can introduce bias and confounding that undermine the accuracy of study results (Astle and Balding 2009; Bhatia et al. 2016; Devlin and Roeder 1999; Hellwege et al. 2017; Kang et al. 2010; Newman et al. 2001; Shibata et al. 2013; Voight and Pritchard 2005; Young et al. 2019). For large-scale biobanks, a substantial portion of study participants may be biologically related; an estimated 32.3% of the individuals in the UK Biobank dataset (Bycroft et al. 2018) have a third-degree or closer relative in the same dataset. Thus, controlling for relatedness can have a major impact on the size and composition of the analysis cohort, and thereby affect the final analysis results. Removal of duplicate individuals across datasets is a special case of detecting relatives, which our work also addresses.

There are several key hurdles to identifying related individuals across datasets. Unlike other analysis tasks that derive aggregate-level insights from the pooled data, such as association tests, finding relatives is an inherently sensitive task, directly operating at the level of individuals. Consequently, most existing approaches for cross-site analysis, e.g., meta-analysis or federated learning, cannot be applied in our setting, as they rely on sharing aggregate-level data between the parties. Furthermore, despite the growing literature on cryptography-based secure computation algorithms for biomedicine (Blatt et al. 2020; Cho et al. 2018; Froelicher et al. 2021b; Cho et al. 2022), which allow joint computation without sharing private data between parties, to our knowledge no practical solution exists for relative detection. This is mainly because standard tools for evaluating kinship require all pairs of individuals between two datasets to be compared (Manichaikul et al. 2010; Purcell and Chang 2023; Conomos et al. 2016) or involve complex combinatorial operations (e.g., string matching) (Gusev et al. 2009; Naseri et al. 2019; Shemirani et al. 2021), which incur an overwhelming cost when implemented using cryptographic operations.

Here we introduce SF-Relate, a scalable and privacy-preserving solution for identifying relatives across distributed datasets, as illustrated in Figure 1. Our novel approach entails each party locally assigning their samples to buckets such that related samples are more likely to be assigned to the same bucket, via *locality sensitive hashing* (LSH) (Indyk and Motwani 1998), and then securely estimating kinship only between samples that end up in the same bucket across parties. We devised a data encoding scheme for LSH aimed at capturing identity-by-descent (IBD) segments in genetic sequences, thus effectively grouping together samples that are likely to be related. Furthermore, we introduce a new strategy for bucket assignment, in which buckets obtained through multiple LSH trials are merged and filtered to obtain a set of size-one buckets (referred to as *micro-buckets*), enabling efficient comparison between parties. We illustrate how our techniques guarantee a high probability of detecting related samples, minimize computational overhead, and ensure that no private information is revealed between the parties. Moreover, to estimate kinship coefficients (Methods) for pairs of samples in the same bucket between parties without sharing data, we introduce a provably end-to-end secure approach that leverages homomorphic encryption, a cryptographic technique allowing for direct computation on encrypted data, combined with efficient distributed computation strategies. SF-Relate keeps each party’s data confidential throughout the computation, revealing only the final output to each party, which includes only the list of their own samples that have at least one relative in another dataset.

**Fig. 1:**
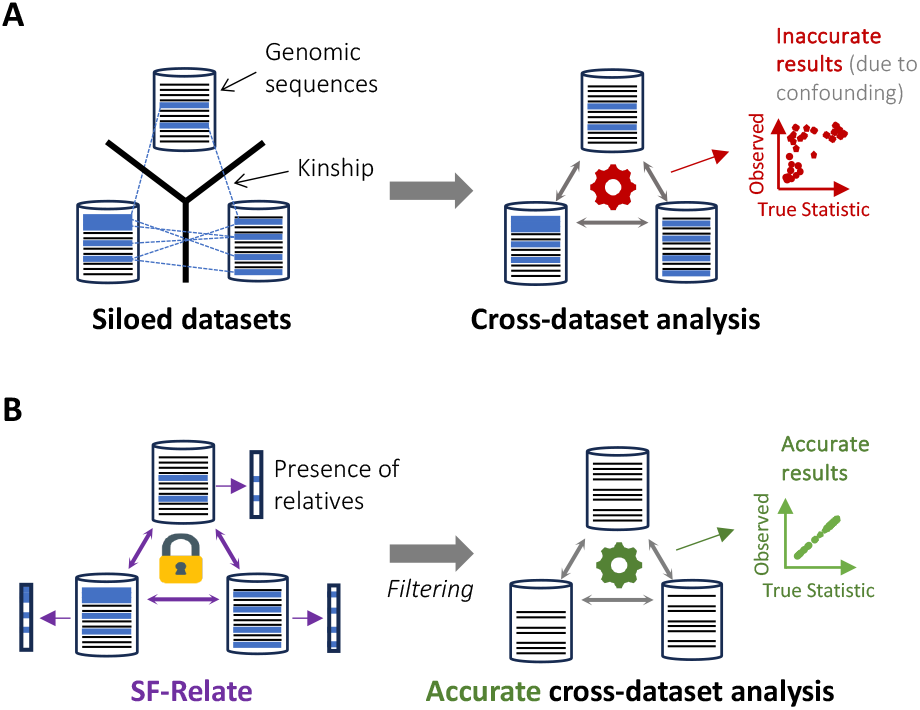
SF-Relate overview. When genetic relatives across datasets cannot be identified due to restricted data sharing, joint studies can suffer from bias and confounding (**A**). SF-Relate allows parties to securely identify and correct for cross-dataset relatives to enhance downstream analysis (**B**).

Our results show that our solution remains practical even for large-scale biobank datasets with a runtime of 14.5 hours to identify the relatives among a pair of datasets consisting of 200K real genomic samples in total. Our privacy-preserving solution enables us to identify more than 97% of the related samples (third-degree or closer) by performing only 0.13% of the computation required by the naive solution in which all pairs of samples are compared between the parties.

## Results

### Overview of SF-Relate

SF-Relate enables multiple parties to detect cross-site relatives in their joint dataset without having to share any sensitive information (Figure 1). The input dataset for each party includes phased haplotype sequences from individuals within that party’s cohort. We consider the parties to be honest-but-curious, meaning that they follow our analysis protocol faithfully but might try to infer information about other parties’ datasets based on what they observe individually during the protocol execution. Based on this model, SF-Relate guarantees end-to-end confidentiality for each party’s input dataset, protecting it from other parties in the protocol. During the protocol, any data exchanged between the parties are encrypted in a manner that requires the participation of *all* parties for decryption, thus ensuring a high level of protection. This approach allows parties to disclose only the information they agree to reveal, such as the final output.

To efficiently scale to large datasets, SF-Relate follows a two-step approach (Methods). In *Step 1: Hashing and Bucketing*, each party locally evaluates a series of hash functions on each individual’s haplotype sequences to assign the individual to buckets across a collection of hash tables, such that related individuals are more likely to be assigned to the same bucket index. For this purpose, we devised a novel encoding scheme that splits and subsamples genotypes into *k-SNPs* (similar to k-mers, but non-contiguous; SNP: single nucleotide polymorphism), such that the similarity between k-SNPs reflects extended runs of identical genotypes, typically indicative of relatedness. We then leverage LSH to derive bucket indices from the k-SNPs. To capitalize on the fact that related samples will likely be assigned to the same buckets multiple times, SF-Relate merges buckets with the same indices across multiple hash tables (produced by different subchromosomes) and then filters every bucket down to a *single* element, thus minimizing the number of costly kinship evaluations. We refer to this as a micro-bucketing strategy. *A somewhat surprising finding of our work is that, despite the extreme level of filtering applied to each bucket during this process, our strategy enables accurate detection of relatives with remarkable efficiency*. At the end of this process, each party obtains a single hash table with size-one buckets, effectively an ordered list of samples.

In *Step 2: Secure Kinship Evaluation*, the parties securely perform element-wise comparisons between their ordered lists of samples from Step 1. Each comparison involves evaluating a standard estimator of the kinship coefficient, KING (Manichaikul et al. 2010). To calculate the estimator without revealing private information between the parties, we employ multiparty homomorphic encryption (MHE). Data encrypted under MHE can be directly used in computation without needing to be decrypted first, and decryption requires the cooperation of all parties. To minimize the computational overhead of MHE, SF-Relate uses sketching techniques on input haplotypes to reduce data dimensionality before performing kinship computations. Furthermore, our protocol is optimized to maximize the use of operations on local, non-encrypted data, which are significantly more efficient than operations on encrypted data. Finally, the encrypted results are compared to relatedness thresholds and aggregated for each individual, providing each party with an indicator that reflects the presence of a close relative in the other dataset.

We detail our algorithms and novel techniques in Methods.

### Datasets and evaluation settings

To evaluate SF-Relate, we obtained three genomic datasets of varying sizes, including a dataset of 20K samples (individuals) with 1M SNPs from the All of Us Research Program (AoU) (All of Us Research Program Investigators 2019) and two datasets from the UK Biobank (UKB) (Bycroft et al. 2018) including 100K and 200K samples, respectively, both with 650K SNPs. The two UKB datasets were uniformly sampled from the full UK Biobank release v3 (*n* = 488, 377), and the AoU dataset comprises the first 20K individuals in the the All of Us release v5 (*n* = 98, 590). We then evenly split each dataset into two parts to emulate a cross-dataset analysis involving two parties. We compute the ground-truth by evaluating all pairwise kinship coefficients using the KING approach ((Manichaikul et al. 2010); see Methods) in plaintexts on a set of ancestry-agnostic SNPs, as in UK Biobank’s pipeline (Bycroft et al. 2018). We provide further details on dataset preparation in Methods.

We evaluate the accuracy of our method in detecting close relatives between two datasets using the standard metrics of *recall* and *precision*. Recall represents the fraction of samples with a close relative in the other dataset (as determined by the baseline KING method given a threshold) that SF-Relate successfully identifies. Precision represents the fraction of samples identified by SF-Relate as having a close relative in the other dataset that actually have such a relationship according to the baseline method. For evaluation of computational costs, we measured the elapsed wall-clock time and the total number of bytes sent from one party to another (given the symmetry of SF-Relate’s computation) for runtime and communication costs, respectively.

All of our experiments were performed using virtual machines (VMs) on the Google Cloud Platform (GCP). This represents a realistic setting where parties use a cloud service provider to access high-performance computing resources that may not be readily available in the local environment. Furthermore, many biobank datasets, including AoU and UKB, are now hosted on cloud-native environments for data analysis. For UKB, we used two VMs (one for each party) with 128 virtual CPUs (vCPUs) and 856 GB of memory (n2-highmem-128) co-located in the same zone in GCP. For AoU, we emulated the two parties in a single VM with 96 vCPUs and 624GB memory due to the constraints of the provided data analysis platform. Supplemental Table S1 summarizes all symbols and default parameters.

### SF-Relate accurately and efficiently detects close relatives between large-scale datasets

We summarize our results on the AoU and UKB datasets in Tables 1 and 2. Across all three datasets (AoU-20K, UKB-100K, and UKB-200K), SF-Relate obtains near-perfect recall and precision (both exceeding 97% in all cases) for detecting the presence of 3rd-degree or closer relationships between two parties. Calculating the recall separately for each relatedness degree from 0th (monozygotic twins) to 3rd, we observe that most missing relationships are for the 3rd degree; SF-Relate finds *all* existing relationships up to the 2nd degree in all three datasets, with the exception of the 2nd degree in UKB-200K, for which it missed 2 out of 1711 individuals with a relative. The recall metric for third-degree relationships remains high—above 94% for all three datasets. Note that the more distant the relationship, the more difficult it is to detect, because the IBD segments become more scattered and reduced in quantity, which in turn results in a lower rate of surviving the filtering step in micro-bucketing. In UKB-200K, we observed that a small fraction (5%) of 3rd-degree relatives, missed by SF-Relate, correspond to those with kinship coefficients near the 4th-degree threshold (Figure 2), suggesting that some of them may not be real 3rd-degree relationships considering the stochastic nature of the kinship estimator.

**Table 1:** SF-Relate achieves near-perfect accuracy for identifying close relatives in UK Biobank and All of Us datasets. Ground-truth relatedness degrees for recall and precision metrics are obtained using the KING method and assigning each sample to the lowest degree of relatedness observed. SF-Relate obtains accurate results while performing only a small fraction of comparisons compared to **all-pairwise** comparison between datasets.

| Dataset | Recall (% , counts) |  |  |  |  | Precision<br>(%, counts) | % of comparisons<br>w.r.t.<br>all-pairwise |
| --- | --- | --- | --- | --- | --- | --- | --- |
|  | Relatedness degree |  |  |  | Overall |  |  |
|  | 0th | 1st | 2nd | 3rd |  |  |  |
| UKB-200K | 100.0%<br>16/16 | 100.0%<br>4702/4702 | 99.8%<br>1709/1711 | 94.9%<br>8475/8925 | 97.0%<br>14902/15354 | 98.5%<br>14902/15129 | 0.13% |
| UKB-100K | 100.0%<br>6/6 | 100.0%<br>1243/1243 | 100.0%<br>404/404 | 95.1%<br>2169/2279 | 97.2%<br>3822/3932 | 98.7%<br>3822/3872 | 0.26% |
| AoU-20K | 100.0%<br>14/14 | 100.0%<br>209/209 | 100.0%<br>93/93 | 94.1%<br>145/154 | 98.0%<br>461/470 | 100.0%<br>461/461 | 1.28% |

**Table 2:** SF-Relate scales efficiently to large datasets. We report the runtime and communication costs for individual steps of SF-Relate described in Methods. The runtime and communication costs for setting up the cryptographic keys are 40.4 s and 1.7 GB, respectively, constant across all experiments. We also show the estimated total costs of running **all-pairwise** comparisons and determining the closest relationship for each individual both using MHE.

| Dataset | SF-Relate |  |  |  |  |  |  |  | All-pairwise |  |
| --- | --- | --- | --- | --- | --- | --- | --- | --- | --- | --- |
|  | Runtime |  |  |  | Communication |  |  |  | Runtime<br>(est. total) | Comm.<br>(est. total) |
|  | Step 1 | Step 2 (MHE) |  | Total | Step 1 | Step 2 (MHE) |  | Total |  |  |
|  |  | Phase 1 | Phase 2 |  |  | Phase 1 | Phase 2 |  |  |  |
| UKB-200K | 1.8 m | 14.0 h | 0.5 h | 14.5 h | — | 46.6 TB | 0.5 GB | 46.6 TB | 1.3 y | 32.5 PB |
| UKB-100K | 49.5 s | 7.05 h | 0.23 h | 7.29 h | — | 23.85 TB | 241.7 MB | 23.85 TB | 112 d | 9.8 PB |
| AoU-20K | 18.6 s | 5.65 h | 0.11 h | 5.79 h | — | 6.2 TB | 77.6 MB | 6.2 TB | 18.8 d | 2.31 PB |

**Fig. 2:**
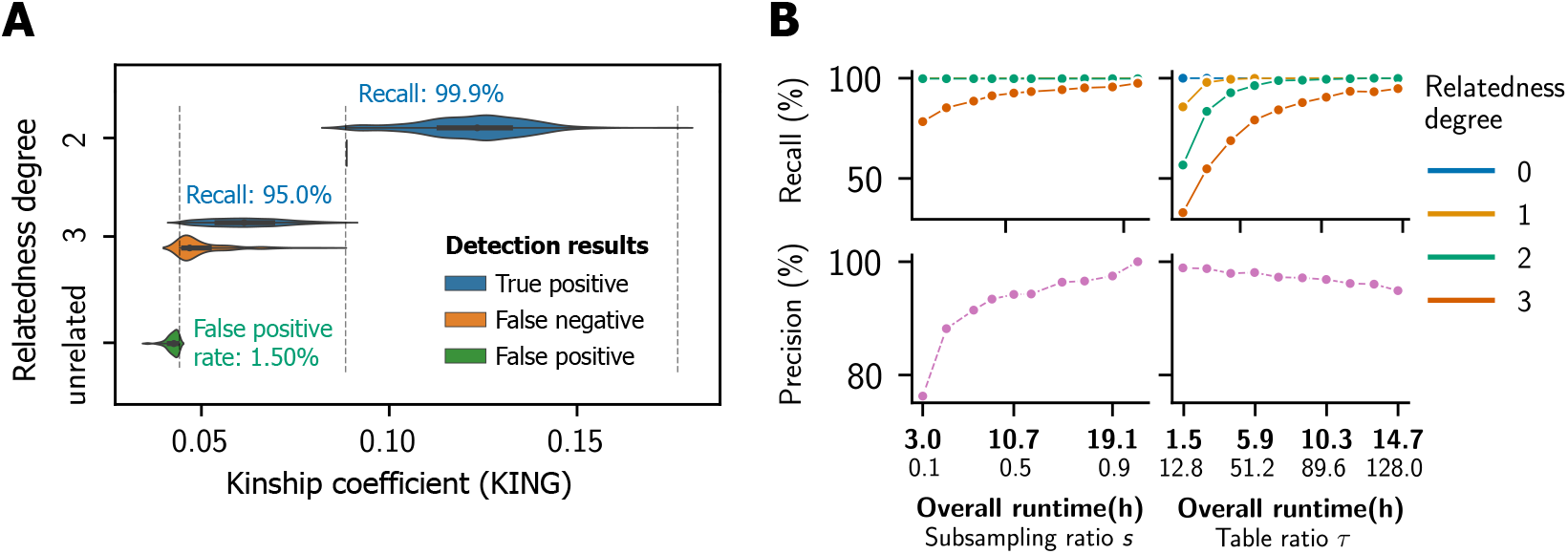
SF-Relate achieves higher accuracy for samples with closer kinship and enables a trade-off between accuracy and runtime. (**A**) We plot the distribution of kinship coefficients (KING) stratified by the (closest) relatedness degree of the relative pairs and by whether they were detected by SF-Relate as related. Mis-classifications by SF-Relate are concentrated around kinship thresholds for different relatedness degrees, indicated by vertical dashed lines. (**B**) We vary the subsampling ratio (*s*) and the table ratio (*τ*) parameters in SF-Relate and report the resulting precision and recall for different relatedness degrees. For precision, only the overall metric is shown for detecting 3rd-degree or closer relatives. By default, *s* = 0.7 and *τ* = 128. There parameters determine the trade-off between the runtime and accuracy of SF-Relate.

Furthermore, SF-Relate consistently achieves high detection accuracy across a variety of populations with distinct ancestry backgrounds. SF-Relate maintains a recall rate exceeding 98% for a dataset comprising individuals of African ancestry (Supplemental Table S2). SF-Relate largely remains effective in multi-ancestry datasets, achieving a recall higher than 80% across all subpopulations (Supplemental Table S3). Ancestry groups with the lowest recall (Indian and Other with 84.4% and 82.4%, respectively) are associated with small sample counts, suggesting that the slight reduction in recall may be due to sampling noise. Taken together, these results demonstrate SF-Relate’s accurate relative detection performance across a range of datasets, which is achieved without revealing any private information between the two parties due to SF-Relate’s use of secure computation techniques when jointly analyzing the two datasets.

Despite the overhead of cryptographic protocols for secure computation, the runtime of SF-Relate remains practical for all three datasets, resulting in 5.8, 7.3, and 14.5 hours of runtime for AoU-20K, UKB-100K, and UKB-200K, respectively. We note that the doubling of runtime from UKB-100K to UKB-200K reflects the linear scaling of SF-Relate in the number of individuals in the dataset, since these datasets were analyzed using the same computing environment, unlike AoU. More precisely, the computational cost of the MHE calculation of pairwise kinship coefficients, which is the main computational bottleneck of SF-Relate, grows linearly in both the number of SNPs after sketching and the size of the hash table. Although both parameters can be adjusted by the user, to maintain accurate performance, these parameters need to be linearly scaled with the total number of SNPs and individuals in the original input dataset, respectively. We provide a systematic evaluation of the runtime scaling of SF-Relate in Supplemental Figure S1. The observed communication costs in the order of tens of terabytes (e.g., 93.2 TB for UKB-200K) are not small, but our results demonstrate that transferring such large amounts of data does not lead to impractical runtimes. In addition, we note that more than 99% of the communication bandwidth is due to the exchange of encrypted hash tables including the sketched haplotypes, which can, in principle, be transferred in a single round of communication. Therefore, we expect the impact of high communication on runtime to be minimal even in a wide-area network (WAN) setting with high communication latency (round-trip delay).

Furthermore, we highlight that without the hashing and bucketing strategy we introduced in SF-Relate, it would not be feasible to securely detect relatives between datasets by all-pairwise computation of the kinship coefficient (**All-pairwise** in Table 2). Even with our efficient MHE implementation of the kinship calculation over the sketched haplotypes, performing all-pairwise comparisons for the UKB-200K dataset is estimated to take 1.3 years based on the same computational setting. On the contrary, SF-Relate obtains practical runtimes by significantly reducing the number of candidate individual pairs to test without compromising accuracy through our novel use of LSH hash tables. Remarkably, SF-Relate makes only 1.28%, 0.26%, and 0.13% of pairwise comparisons compared to the total number of individual pairs between the two datasets for AoU-20K, UKB-100K, and UKB-200K, respectively (Table 1). This drastically reduces not only the runtime but also the communication costs; e.g., our MHE computation would require 65 PB of communication without our hashing techniques (Table 2).

### Navigating SF-Relate’s accuracy-runtime tradeoffs and parametrization

The runtime and accuracy of SF-Relate are primarily influenced by the hash table size *N* = *τ* · *n*, determined by the dataset size *n* and table ratio (*τ*; Methods), and the sample size for comparison, determined by the subsampling ratio (*s*) and the number of SNPs (Methods). As shown in Figure 2, increasing *s* improves the overall recall and precision, while increasing *τ* enables the detection of more distant relationships, also increasing overall recall. However, the runtime depends linearly on both *s* and *τ*, highlighting the trade-off between SF-Relate’s accuracy and runtime. We expect the optimal trade-off to depend on the application setting.

For example, if users want to focus on identifying relatives up to the 2nd degree within the UKB-200K dataset, they could set the table ratio *τ* to 64 and the subsampling rate *s* to 0.7, instead of *τ* = 128 and *s* = 0.7 in our experiments (Methods). This results in a two-fold improvement with respect to our experiments due to halving of the hash table size. Even in this scenario, users would maintain an effective detection rate of over 95% for individuals with relationships closer than the 2nd degree (Figure 2B). Alternatively, if users want to achieve perfect accuracy, they can increase *s* and *τ*. Increasing *s* from 0.7 to 1 (i.e., no sketching), improved SF-Relate’s overall recall on UKB-200K from 97.0% to 98.7% and precision from 98.5% to 99.9% (Table 1 and Supplemental Figure S2). The runtime increased from 14 to 21 hours. Furthermore, by doubling the table ratio *τ*, SF-Relate achieves perfect accuracy for relations up to the 3rd degree, while doubling its runtime. Overall, SF-Relate’s recall remains consistently high, above 95%, across a wide range of parameters, and only starts to decrease when the parameters significantly deviate from the default setting (Supplementary Table S4).

To choose suitable values for *s* and *τ* in practice, we recommend that users first determine the farthest relationships they wish to detect and an acceptable level of recall. Using Figure 2, they can then determine the required *s* and hash table size *N* = *τn*. The expected runtime can be estimated by considering the linear relationship between these parameters and SF-Relate’s runtime. To select the *Hashing and Bucketing* parameters, the users may initially opt for our recommended parameters (Supplemental Table S1), which consistently achieve accurate and efficient performance across all our datasets. Optionally, users can conduct a local assessment using their own dataset to verify and fine-tune the chosen parameters, which involves examining the number of shared genomic segments between local relatives and their matching probabilities (as depicted in Supplemental Figure S3) and estimating the detection rate among local relatives.

### SF-Relate’s IBD-based hashing strategy improves detection accuracy over the KING estimator

SF-Relate leverages a secure implementation of the KING formula (Manichaikul et al. 2010) to estimate relatedness (Methods). Nevertheless, we observed that SF-Relate can sometimes lead to even more accurate identification of relatives than KING. This is due to SF-Relate’s IBD-based approach to bucketing the samples, which helps filter out spurious pairs of samples that are identified by KING as being related, which in fact do not share any long IBD segments. We confirmed this in our comparisons with PC-Relate (Conomos et al. 2016) and RAFFI (Naseri et al. 2021), recent methods for kinship detection designed to improve upon KING’s accuracy by correcting for population structure and incorporating IBD segment detection, respectively. We evaluated all methods on a subset of 20K samples from the UKB-200K, distributed across two parties. As expected, they produce highly similar results for up to the 3rd degree relatives (Figure 3, Supplemental Table S5, and Supplemental Figure S4). However, for detecting 4th-degree relatives, standard KING erroneously identified numerous individuals as being related due to the presence of an outlier sample that is detected as related to *thousands* of samples in the other dataset; SF-Relate, akin to the more advanced tools PC-Relate and RAFFI, successfully avoids these errors. Similar outlier-related issues regarding KING have been noted in UK Biobank’s official report on relatedness inference (Supplementary in (Bycroft et al. 2018)). In Figure 3, we visualize sequence similarity between four pairs of haplotypes involving the outlier sample, compared with typical 4th-degree relative pairs identified by PC-Relate. We observe light yellow bands of high sequence similarity regions exclusively in PC-Relate’s pairs, which signifies real IBD segments. This suggests that SF-Relate’s bucketing approach based on IBD segments can effectively distinguish outlier pairs from real ones, thus leading to more accurate detection of relatives.

**Fig. 3:**
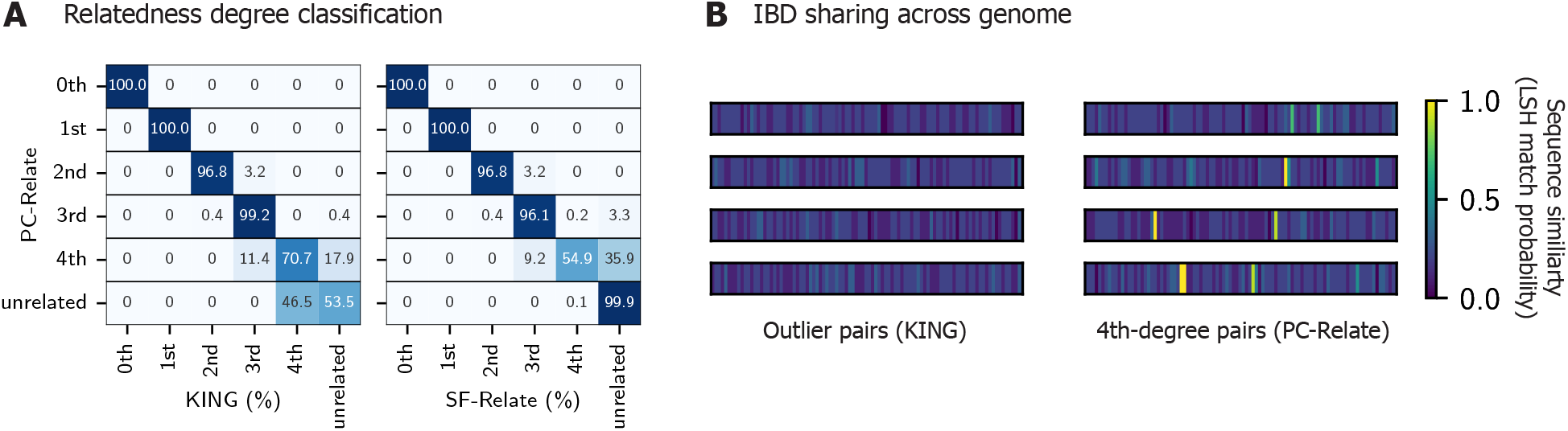
SF-Relate excludes spurious 4th-degree relatives detected by KING. (**A**) We show confusion matrices assessing the relatedness classification accuracy of KING (left) and SF-Relate (right), comparing with the output of PC-Relate as the ground-truth. SF-Relate is performed in plaintext (i.e., without MHE), focusing on the evaluation of the bucket assignment. Unlike SF-Relate, KING classifies many unrelated samples as 4-th degree relatives. Most of these pairs involve the same outlier sample, which has many spurious relationships. (**B**) We verify that pairs involving the outlier do not exhibit IBD sharing patterns (left), evident in 4th-degree pairs from PC-Relate (right). Four example pairs are shown for both cases. For each pair, we compute the Hamming similarity between the two samples of genomic segments across the genome. Bright yellow bands represent likely IBD segments. The locations of the bands are randomly permuted to obscure their positions.

### SF-Relate supports alternative output settings

In SF-Relate’s default setting, each party learns only whether each local individual has a close relative within the joint dataset. SF-Relate offers an option to output kinship computation results with more detailed granularity, summarizing them at various levels to address a range of analysis needs. Alternative outputs include the closest relatedness degree for each individual, the maximum kinship coefficient for each individual (discretized), and the full list of computed kinship coefficients. The SF-Relate output remains accurate in all settings: The individual kinship coefficients computed by SF-Relate exhibit a small average absolute error of 5.8 × 10^−4^ when compared to KING (Supplemental Figure S5). The closest degree reported for each individual matched with the ground truth for 99.9% of the individuals (Supplemental Figure S6). Calculating the maximum kinship coefficient, discretized using the smallest bin width of 0.016, SF-Relate accurately assigned more than 85% of the samples to the correct bins, and more than 99.9% were within one bin of the true output (Supplemental Figure S6).

### A case study: SF-Relate reduces false positives in genome-wide association studies

We show that SF-Relate can be used to improve the accuracy of downstream studies without requiring the sharing of sensitive information. We illustrate this using the GWAS workflow and demonstrate SF-Relate’s effectiveness in mitigating confounding from cryptic relatedness by enabling parties to detect and remove relatives from their joint dataset prior to conducting the GWAS. We simulate a multi-site GWAS using a subset of 100K samples from white British participants in UKB, distributed geographically among six parties (Methods and Supplemental Table S6). 50% of these samples have at least one relative of 3rd-degree or less in the dataset. We then simulate 100 phenotypes and perform a linear regression-based GWAS with the top principal component as a covariate (Methods). On average, using a nominal significance cut-off *p*-value of 0.05, SF-Relate removes 2.60% of the falsely identified loci (false positives) with a drop in false positive rate (FPR) from 5.14% to 5.01%, compared to when relatives are not removed from the dataset (Figure 4). When parties independently remove local relatives, 18% of the remaining samples still have relatives in the joint dataset, and 2.03% of false positives are removed with a FPR drop from 5.14% to 5.04%. The one-sided Mann-Whitney U test *p*-value that SF-Relate produces lower FPRs across all phenotypes compared to the local removal of relatives is 1.25 × 10^−5^. Thus, SF-Relate significantly mitigates confounding, producing a FPR near the nominal cutoff *p*-value of 0.05, comparable to centrally-coordinated sample removal (Figure 4).

**Fig. 4:**
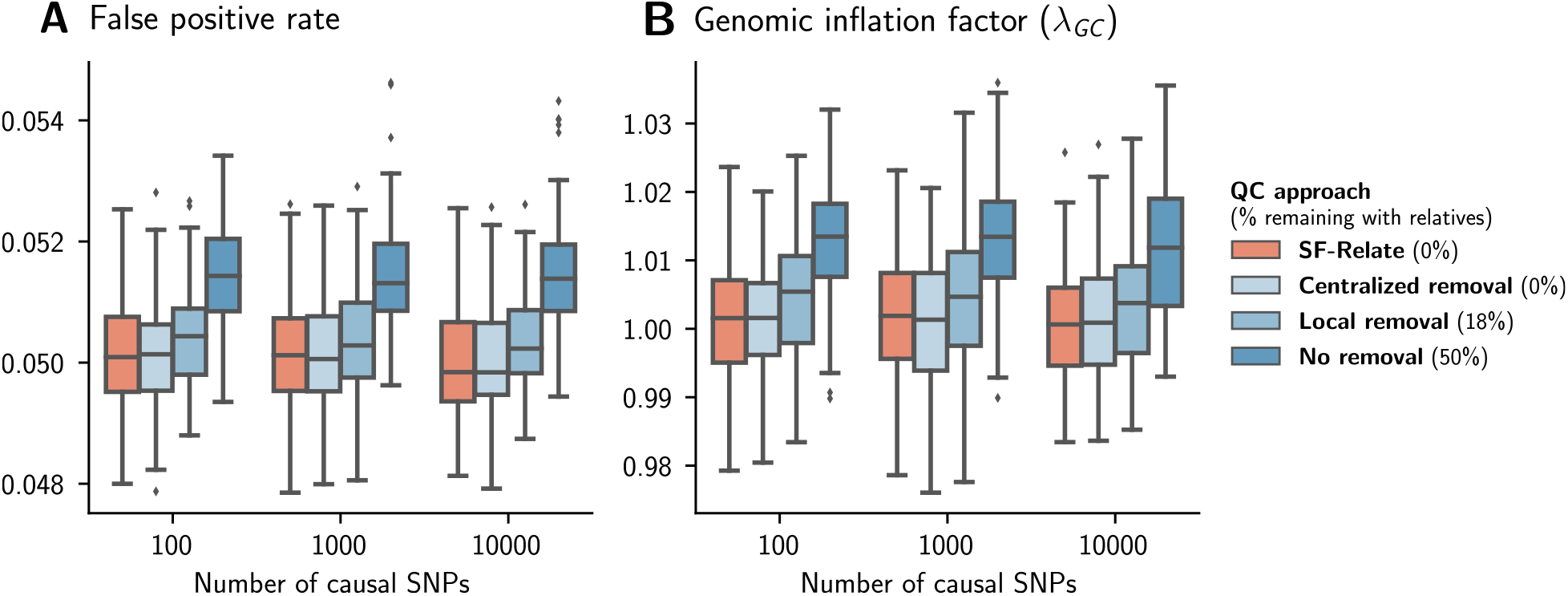
SF-Relate reduces false positives in multi-site GWAS. We vary the number of causal SNPs used in simulating the phenotypes, and compare four quality control (QC) approaches for excluding related individuals: (i) *Centralized removal* (non-private): all relatives are removed from the pooled dataset, (ii) *SF-Relate*: relatives are removed using our secure approach, (iii) *Local removal*: each party filters relatives from its local dataset independently, and (iv) *No removal*: no relatives removed. Initially, 50% of samples have relatives, and *Local removal* results in 18% of remaining samples still having a relative in the joint dataset. We plot the fraction of significant loci (*p*-value *<* 0.05) on even numbered chromosomes in that are designed to be non-causal in the simulation (**A**), and the genome inflation factor *λ*_*GC*_ in (**B**). The filled boxes represent interquantile ranges of statistics across 100 simulated phenotypes. While local removal of relatives help reducing the confounding to some extent, SF-Relate significantly mitigates confounding, comparable to centrally-coordinated sample removal.

## Discussion

We presented SF-Relate, a secure federated algorithm for identifying close relatives between isolated genomic datasets. Using a novel strategy for hashing and bucketing individuals to capture shared IBD segments between relatives, SF-Relate achieves near-perfect detection while maintaining a practical runtime (i.e., less than a day) even on a large dataset including 200K individuals. To the best of our knowledge, our work is the first to demonstrate secure relative detection at scale while also ensuring a strong, formal notion of privacy for each input dataset. We expect SF-Relate to be a useful tool for the growing networks of collaborating institutions, which currently lack the tools to jointly perform a variety of genetic analyses without sharing data. To facilitate the use of SF-Relate, we provide automated deployment workflows on the sfkit web server (Mendelsohn et al. 2023), which streamlines the collaborative execution of a range of secure federated tools such as SF-Relate.

There are several directions that we would like to pursue in future research. First, while SF-Relate identifies relatives based on the standard KING-robust estimator (Manichaikul et al. 2010), there are other approaches that may provide more robust estimation, especially for more distant relatives beyond the third degree, in terms of both correcting for population structure (e.g., PC-Relate (Conomos et al. 2016)) and detecting IBD segments to allow a more direct calculation of the proportion of IBD sharing (e.g., RAFFI (Naseri et al. 2021)). Although we have demonstrated that our method can often mirror the behavior of these advanced methods (Figure 3; Supplemental Table S5 and Supplemental Figure S4), directly implementing these approaches may be more effective for identifying distant relatives. Integrating our approach with a recently proposed secure federated algorithm for principal component analysis (Froelicher et al. 2023), may help to address the former. For the latter, we posit that an extension of our hashing strategy to quantify the rate of collision, which represents the sharing of a short IBD segment between individuals, may be possible.

Extending SF-Relate to accommodate a broader range of scenarios represents another key direction of future work. For scenarios involving more than two parties, developing a more efficient strategy than the straightforward all-pairwise execution of SF-Relate would be beneficial. Additionally, enabling the detection of relatives for a single query individual within a large (distributed) database would be useful for services that help individuals find lost biological relatives (e.g., MyHeritage (2023)). There is also a need to facilitate similarity computations for other data types, including medical records. In any of these scenarios, it would be meaningful to further explore potential information leakage in the output and devise strategies to mitigate any remaining risk. Overall, our work offers technical insights that are broadly applicable to discovering relations across siloed datasets.

## Methods

### Overview of the Cross-Dataset Kinship Estimation Problem

We consider the setting in which multiple parties (e.g., researchers from different institutions) wish to identify genetically related individuals between their private datasets while protecting the confidentiality of their data (**Fig. 1**). The goal of the parties is to use this information to facilitate downstream collaborative analysis by excluding duplicate or related individuals from the analysis to minimize bias in statistical analyses. For simplicity, we focus on the setting with two parties, each holding a dataset including phased haplotypes of *n* individuals over *m* genetic variants, such as single nucleotide polymorphisms (SNPs). The desired output for each party is a list of individuals in their private dataset who have at least one “close” relative in the other party’s dataset with respect to some relatedness threshold. In our work, we mainly consider the detection of third-degree or closer relatives, which are the closest relations most commonly used in genomic studies and data releases (e.g., UK Biobank (Bycroft et al. 2018)). We assume each input dataset to be locally phased by each party (i.e., each individual’s genome is represented as two haplotype sequences), which is crucial for capturing identity-by-descent (IBD) sharing patterns, as we describe later. We further consider a threat model where the parties are *honest-but-curious*; i.e., they faithfully follow the protocol as specified, but might attempt to infer private information about the other parties’ datasets based on the data observed during the process. Given this model, we aim to provide formal privacy guarantees for each party’s input dataset, ensuring that no information is revealed to other parties except for what can be gleaned from each party’s respective output and global parameters of the problem (e.g., dataset dimensions and security parameters).

### Estimation of kinship coefficients

The *kinship coefficient ϕ* between a pair of individuals is defined as the probability that a pair of randomly sampled alleles is identical by descent (IBD), i.e., when the pair of alleles is identical due to genetic inheritance rather than by chance. For example, since human genomes are diploid, a direct descendant inherits exactly one set of chromosomes from each parent; in this case, the kinship coefficient between an individual and his or her parent is, in principle, 0.5 · 0.5 = 0.25.

Existing approaches for estimating the kinship coefficients typically fall into one of two classes: *distance-based methods*, which use a notion of distance between two genotype vectors that often incorporate information about minor allele frequencies (MAF) (Conomos et al. 2016; Manichaikul et al. 2010); and *IBD-segment-based methods*, which first identify long shared segments between individuals that are likely due to IBD (Nait Saada et al. 2020; Naseri et al. 2019; Shemirani et al. 2021) and estimate kinship based on the extent of these shared segments. In a benchmarking study, Ramstetter et al. (2017) showed that both approaches achieve high accuracy for up to third-degree relationships, while agreement becomes weaker for more distant relationships, which we also observe between PC-Relate and KING (Figure 3). Although IBD-segment-based methods generally offer a more accurate estimation of kinship by analyzing IBD sharing patterns, distance-based approaches represent an efficient alternative that does not involve costly string matching, which often leads to substantially higher runtime (e.g., days vs. minutes (Ramstetter et al. 2017)). As a result, distance-based methods have been more commonly applied to large datasets (Bycroft et al. 2018). Recently proposed methods (e.g., RaPID (Naseri et al. 2019)) introduce hashing techniques to improve the scalability of IBD segment detection, which in some cases exceed the efficiency of distance-based methods due to the quadratic scaling of pairwise distance calculation. However, IBD segment finding methods are still combinatorial in nature and cannot be efficiently implemented using existing secure computation techniques. In our work, we adopt a distance-based approach to minimize the computational overhead associated with secure computation but simultaneously exploit IBD sharing patterns to significantly improve the scalability of our approach.

Our work addresses the problem of applying the widely adopted distance-based method for kinship estimation, the KING-robust estimator (referred to as KING in what follows for simplicity) (Manichaikul et al. 2010), to find relationships between two datasets. This method is implemented in several standard genomic analysis toolkits, such as Hail (Hail 2023) and PLINK (Purcell and Chang 2023), and recently the UK Biobank released relatedness data for individuals in the dataset using this estimator (Bycroft et al. 2018). KING estimates the kinship coefficient between two genotype vectors **x** and **y** ∈ {0, 1, 2}^*m*^ using the following formula:

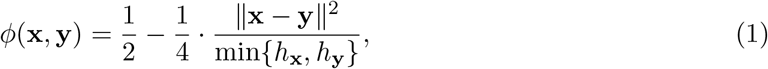

where each element in **x** and **y** represents the minor allele dosage of a genetic variant, and *h*_**x**_ and *h*_**y**_ represent the fraction of heterozygous loci in each vector (i.e., the heterozygosity of the individual).

### Existing cross-dataset approaches and their limitations

Wang et al. (Wang et al. 2022) proposed a homomorphic encryption method for identifying genetic relationships across parties, but their approach requires kinship computation for *all pairs* of samples, which does not scale to large datasets. Other previous approaches (Dervishi et al. 2023; Glusman et al. 2017; Hormozdiari et al. 2014; Robinson and Glusman 2018) rely on sharing a limited amount of processed data between parties to find related samples, which sacrifices both privacy and accuracy to some extent. For instance, Dervishi et al. (Dervishi et al. 2023) introduced a solution in which the parties reveal a subset of SNPs in a shuffled order for their respective samples to estimate the kinship coefficients. Robinson and Glusman (2018) and Glusman et al. (2017) proposed to compare “fingerprints” obtained by applying a random projection to genomic samples to infer relatedness. He et al. (2014) and Hormozdiari et al. (2014) used error-correcting codes and fuzzy encryption to compare genotype vectors such that the comparison result can be decoded only if the two vectors are similar enough. These solutions require comparison between all pairs of samples between datasets, and the processing of genotype vectors into limited representations that can be shared leads to loss of precision.

In a recent competition organized by the iDASH Workshop 2023 (iDASH 2023), identifying the presence of relatives in encrypted datasets was posed as one of the challenge tasks. The challenge considered a setting that is different from our work. It involved a client-server scenario with small datasets (e.g., 2K individuals and 16K variants) and restricted any data preprocessing for evaluation purposes, thus limiting the applicability of the proposed solutions in a real use-case scenario. The best-performing solutions to the challenge generally followed a similar approach adapting the work of Homer et al. (2008). In this approach, the presence of a relative in a dataset is inferred based on a statistic evaluating whether the individual’s genotype vector is closer to the allele frequencies in the dataset than to background frequencies computed in a reference population. We show in Supplemental Figure S7 that computing this statistic securely does not lead to accurate results in realistic settings, involving complete genomes and a large number of individuals (e.g., tens of thousands or more).

### Our novel approach to securely detecting relatives across large-scale and distributed datasets

We developed SF-Relate to allow secure and efficient detection of relatives between large-scale datasets by drastically reducing the number of kinship computations, from quadratic to linear, while preserving accuracy. For a graphical overview of SF-Relate’s workflow, see Supplemental Figure S8. To achieve this solution, SF-Relate draws ideas from both *distance-based* and *IBD-segment-based* kinship estimation methods. It first identifies pairs of samples to be compared between parties using a locality sensitive hash (LSH) function (Indyk and Motwani 1998), which we adapt to ensure that both (or all) parties assign individuals with shared IBD segments to the same bucket with a higher probability than for unrelated individuals. SF-Relate sets the capacity of each bucket to one, discarding duplicate hits, which we refer to as a *micro-bucketing* strategy. As we demonstrate, this novel approach is key to minimizing the number of comparisons while maintaining accuracy. Next, SF-Relate securely estimates and thresholds the kinship coefficients between pairs of samples in buckets with the same index across parties then aggregates the results per sample using our secure implementation of a distance-based kinship estimator (KING-robust (KING 2023)). To perform these operations while keeping each party’s data confidential from other parties, we develop efficient two-party computation protocols based on multiparty homomorphic encryption (MHE) techniques (Mouchet et al. 2021). We additionally incorporate sketching techniques to further reduce the computational cost of MHE computations. In the following, we provide details of each step of our algorithm.

### Step 1: Hashing and bucketing

In this step, each party locally evaluates a series of hash functions on each individual’s haplotype sequences to assign the individual to buckets across a collection of hash tables, such that related individuals are more likely to be assigned to the same bucket index. Only individuals in the same bucket across parties are compared in a later step.

#### Locality sensitive hashing to capture IBD segments

SF-Relate assigns individuals to buckets using a locality sensitivity hash (LSH) (Indyk and Motwani 1998). LSH functions map similar items to the same value more frequently based on a similarity notion. Specifically, the Hamming LSH, which relies on the Hamming similarity (defined as the number of equal coordinates between vectors), projects a vector onto one random coordinate and uses its value as the output. In our setting, applying the Hamming LSH (or other LSH methods, such as MinHash (Broder 1997)) directly to each sample, encoded as a genotype vector, for bucket assignment would not work in practice. This is because the difference between related and unrelated individuals, with respect to the distance between samples’ encodings, is too small for LSH to distinguish; the average relative Hamming distance between third-degree relatives and unrelated individuals in UK Biobank are 22% and 23%, respectively. Hence, SF-Relate applies an encoding scheme that results in highly similar Hamming vectors for related samples. This encoding captures the biological signal of IBD distributions, and can be seen as a variant of the encoding in (Shemirani et al. 2021). It applies LSH on split chromosomes, exploiting the key insight that IBD segments are unevenly distributed on the genome. The subchromosomes are further divided into short string of genotypes (similar to k-mers), to extract long runs of identical genotypes that are unlikely by chance.

The first three steps of our hashing approach encode IBD segments: (1) *Splitting*: Haplotypes are divided into genomic segments of fixed-length in genetic distance (centi-Morgans). (2) *Subsampling*: Each segment is randomly projected down to a fixed number of SNPs, where the SNPs are sampled with probability proportional to their minor-allele frequency (MAF). This reduces the impact of genotyping errors and rare variants on hashing and unifies the SNP density across different segments and datasets, inspired by Naseri et al. (2019). (3) *K-merization*: Subsampled genotypes for several contiguous segments are concatenated to form a k-SNP (akin to *k*-shingles in (Shemirani et al. 2021)), i.e., a sequence of genotypes for *k* SNPs, which helps to identify matches of long genomic segments. Through these steps, we encode each sample as a list of subchromosome k-SNP vectors. In fact, we confirmed that related pairs share significantly more Hamming-similar subchromosomes under this encoding scheme (Supplemental Figure S3).

The final step of hashing applies LSH on the subchromosome vectors to obtain the actual bucket index. To utilize the raw probability gap between non-IBD segments and IBD-segments produced by LSH (such as the gap between 0.5 and 1.0 in Supplemental Figure S3) we need to amplify it. For this, we choose a concatenation parameter *ℓ* and define the bucket index as the FNV-1 hash (Eastlake et al. 2019) on the concatenation of the outputs of *ℓ* independent Hamming LSH applied to the subchromosome vector. This in turn boosts the gap by raising it to the power of *ℓ*. An alternative (as in (Shemirani et al. 2021)) would be to apply the LSH function MinHash (Broder 1997), which captures the set-based Jaccard similarity. Nevertheless, Hamming similarity is more natural for IBD detection, as IBD segments are by definition matching k-SNPs at the same genetic positions, while the Jaccard similarity discards positional information of the two sets of k-SNPs. Indeed, Hamming similarity detects more highly similar subchromosomes (Supplemental Figure S9). The final output of this procedure is a list of hash tables, each consisting of buckets storing sample IDs.

#### Micro-bucketing strategy

To prevent leakage of private information, the parties need to compare the samples in corresponding buckets without exposing any additional information about their datasets, such as the distribution of non-empty buckets and their sizes. This requires that the buckets created in the previous step be padded to a fixed size by adding dummy samples. However, to keep the sample identities hidden, all pairs of samples in a bucket between the parties need to be compared, which leads to both quadratic scaling of comparisons with the bucket size and a large amount of wasted computation involving dummy samples. To address this issue, SF-Relate introduces a *micro-bucketing* strategy, which merges buckets across multiple hash tables (produced by different subchromosomes) and then filters every bucket down to a *single* element. Dummy samples are added only at the end to pad empty bucket to size one. This effectively transforms the parties’ local bucket assignments into an ordered list of samples to be securely compared against the corresponding list obtained by the other party in an element-wise fashion. This approach avoids the quadratic scaling while minimizing the addition of dummy samples due to the merging of buckets (i.e., a bucket is filled if at least one sample is assigned to it in one of the hash tables). Despite the extreme level of filtering applied to each bucket during this process, our strategy enables accurate detection of relatives with remarkable efficiency.

SF-Relate chooses a **table size parameter** *N*, and a **bucket size parameter** *C*. It aligns the different hash tables and merge buckets with the same remainder modulo *N* into one. This ensures the number of buckets is at most *N*. In practice, on a dataset with *n* local samples, *N* is determined by first choosing a **table ratio** *τ*, and letting *N* = *τn*. (A table of the main symbols and their default values is provided in Supplemental Table S1.) After this merge, buckets with more than *C* samples are filtered until *C* samples remain. In this filtering step, samples from buckets built by smaller subchromosome index are given higher preference, but otherwise a uniformly random filtering is performed. The preference towards smaller subchromosome indices is to ensure samples in corresponding buckets in hash tables more likely originates from the same hash table. The parties then repeat the entire hashing step locally *L* times (each with new randomness) until 99% of the buckets are full. Only at the end, dummy samples are inserted to ensure a constant bucket size.

On the realistic datasets UKB-200K, we evaluate the best bucket capacity *C*, by keeping the number of comparisons *NC*^2^ fixed and checking the fraction of related pairs that appear in at least one corresponding bucket after micro-bucketing. Surprisingly, the minimal capacity *C* = 1 gives the highest recall. This is because related samples share many rare IBD segments (many of which is unique) that cause them to end up in small-sized bucket (Supplemental Figure S3). When this happens, both samples remain in the corresponding bucket with high probability (as there is no competition within that bucket). The average fraction of shared unqiue IBD segments is not high for 3rd degree (20/352), but given that our goal is to use the list of buckets to trigger subsequent kinship computations (which operates on the entire genome), and not identifying all IBD segments between pairs (unlike (Shemirani et al. 2021; Naseri et al. 2019)), as long as one of the many small-sized buckets due to rare IBD segments survive micro-bucketing, the pair of relatives would be discovered. This explains the effectiveness of micro-bucketing when restricted micro capacities *C*. In fact, for the rest of the paper, we always set *C* = 1.

In sum, each party in SF-Relate obtains a single hash table with *N* buckets (each with *C* = 1 samples), and micro-bucketing ensures the hash tables are highly utilized. The parties hence perform *C*^2^*N* = *N* secure kinship computation in *Step 2: Secure kinship evaluation*.

### Step 2: Secure kinship evaluation

In this step, the parties perform element-wise comparisons between their ordered list of samples (representing elements in a merged hash table with size-one buckets), obtained in Step 1. They first jointly evaluate the kinship coefficient for each pair (**MHE-Phase 1**), before aggregating the results to obtain an indicator for each individual reflecting the presence of a close relative in the other dataset (**MHE-Phase 2**).

#### Review of multiparty homomorphic encryption (MHE)

To compute on encrypted data in a secure manner, SF-Relate builds upon multiparty homomorphic encryption (MHE) (Mouchet et al. 2021; Froelicher et al. 2021a), extending the CKKS scheme (Cheon et al. 2017). CKKS encodes a vector of real number values in a single ciphertext and is well-suited for calculations where a small amount of noise can be tolerated. Like other HE schemes, it provides operations for addition, multiplication, and rotation (i.e., permutation of elements in a vector) of encrypted values in a ciphertext while providing the single instruction, multiple data (SIMD) property. Operations involving plaintext (unencrypted) data are substantially more computationally efficient; e.g., ciphertext-plaintext multiplication is seven times faster than ciphertext-ciphertext multiplication based on our parameter setting. MHE extends the CKKS scheme to the setting with multiple parties, by secret-sharing the decryption key and constructing a shared collective encryption key. Any party can encrypt data and perform homomorphic computations locally, but decryption can be performed only if all parties cooperate. Moreover, in the multiparty setting, some heavy cryptographic operations, e.g., the bootstrapping which is required to “refresh” a ciphertext after a certain number of multiplications, can be replaced by lightweight interactive protocols (Mouchet et al. 2021; Froelicher et al. 2021a). In SF-Relate, all exchanged data are encrypted under the collective encryption key and only the final result can be decrypted with the cooperation of all parties.

In the following, we describe our efficient MHE-based secure computation protocols for kinship evaluation between two parties (MHE-Phases 1 and 2) based on the bucketed samples from Step 1. We exploit key properties of MHE to minimize the cryptographic overhead of our protocols by maximizing the use of locally available plaintext data and balancing the workload between the parties. Although we focus on the two-party setting, our protocols naturally extend to settings with more than two parties, since we can execute the protocols between all pairs of parties and then aggregate the results.

#### MHE-Phase 1: Secure computation of kinship coefficients

Given a list of *N* samples on both sides, where each sample is associated with a vector of *M* SNPs, the parties collaborate to calculate kinship coefficients for the *N* pairs of samples between them and compare each to a threshold. The desired comparison test given a threshold *θ* is 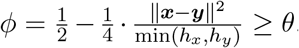. For efficient evaluation under encryption, we can rewrite it as (2 − 4*θ*) min(*h*_*x*_, *h*_*y*_) −∥***x*** − ***y***∥^2^ ≥ 0, thus avoiding the division operation. The comparison test passes when both *h*_*x*_ and *h*_*y*_ satisfies it, so we compute 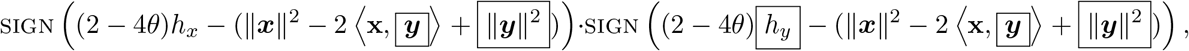 which evaluates to 1 if the coefficient is above the threshold and 0 otherwise. Boxed terms represent encrypted data, and 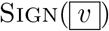 is the indicator function for *υ* ≥ 0. Note that we assign the evaluation of this expression to the party that holds ***x*** (Party 1) and have the other party (Party 2) transfer the encrypted ***y*** for computation. This allows most operations to be performed efficiently using the plaintext ***x***. We observe that *h*_*y*_ (the number of heterozygous in the genotype) and ∥***y***∥^2^ can be computed locally by Party 2 before encryption. Hence, the most expensive operation is the inner product 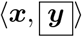 which requires a plaintext-ciphertext multiplication between vectors of size *M* and a summation of elements of the resulting vector. We use a polynomial approximation of the sign function to homomorphically evaluate Sign(·). The cost of this function, despite requiring homomorphic evaluation of a high-degree polynomial, is dwarfed by the cost of computing the inner products. Finally, in addition to the SIMD property of MHE operations, we process batches of coefficients in parallel and evenly distribute the workload between parties by alternating their roles across batches with respect to who holds the plaintext vector ***x***. Our protocol is provided in Supplemental Note S1.

#### MHE-Phase 2: Secure aggregation of results for individual samples

Next, the parties aggregate the comparison results for each individual to compute a single binary indicator representing the presence of a relative. For this, they first perform a linear scan over the comparison results, which selects the results corresponding to the same individual and masks the rest. The selected results are then accumulated, after which another sign test is performed to obtain a binary value as desired, hiding the number of identified relationships as a result. At the end of the protocol, the parties decrypt the vectors and each obtain a list of indicators, determining whether each sample has at least one close relative in the other dataset; this is the only information that is revealed to each party. Note that the complexity of this step depends only on the number of comparisons and individuals, while MHE-Phase 1 also scales with the number of SNPs, which is typically large (e.g., *>*500K). We provide our protocol for this step in Supplemental Note S1.

#### Accelerating kinship computation using sketching

To further reduce the cost of secure kinship evaluation, SF-Relate first reduces the size of each sample through *sketching*. In particular, given the **subsampling ratio** parameter 0 *< s* ≤ 1, SF-Relate randomly chooses a *s* fraction of SNPs to use for kinship evaluation. This provides a natural approximation for the KING kinship estimator, which includes squared Euclidean distance between two genotype vectors that can be estimated using a random subset of coordinates in an unbiased manner (see Supplemental Figure S10). Our results show that this approach enables a meaningful trade-off between accuracy and efficiency; a minor loss in precision introduced by sketching allows us to obtain a substantial reduction in computational cost while maintaining near-perfect detection accuracy.

#### Alternative output modes

By default, SF-Relate computes a list of indicators representing whether each sample has at least one close relative in the other datasets. SF-Relate also supports securely computing other types of output, including the closest relatedness degree for each individual, the maximum kinship for each individual (discretized), and the full list of computed kinship coefficients.

For outputting all kinship coefficients, we replace the final sign test computations with the computation of the kinship coefficients in **MHE-Phase 1** (Supplemental Note S1). That is, the parties homomorphically compute 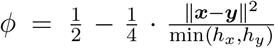. To speed up the computation, they precompute the values 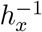 and 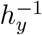 locally in plaintext, which allows replacing the division with a multiplication by 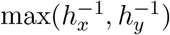, which can be more efficienctly computed.

For both the closest relatedness degree and the maximum kinship coefficient, SF-Relate computes multiple comparison tests with respect to a series of kinship thresholds, then executes **MHE-Phase 2** in parallel to accumulate the resulting comparison results. The decryption of these results reveals the largest threshold at which the comparison test succeeded. Based on this information, the parties can determine the closest degree or the maximum kinship as desired.

Note that, in any of these settings, the final results can also be kept in encrypted form and utilized in subsequent analysis steps without revealing the results of relative detection. Additionally, the complexity of **MHE-Phase 1** is constant across all output modes with *N* plaintext-ciphertext multiplications between vectors of length *M*, whereas the complexity of **MHE-Phase 2** increases linearly with the number of thresholds *t*. The default setting of SF-Relate corresponds to *t* = 1, while the version that reveals all coefficients corresponds to *t* = 0.

### Dataset preprocessing

We utilize three datasets sampled from prominent genomic data consortia: UK Biobank (UKB) and All of Us (AoU). Specifically, we extract two datasets from UKB, one comprising 100K samples (UKB-100K) and the other comprising 200K samples (UKB-200K). In both cases, we randomly split the datasets among two sites. From AoU, we extracted a dataset of 20K samples. Due to the smaller size of this dataset and to avoid an highly imbalanced distribution, we first split the individuals with close relations on the two sites, and then randomly split the set of unrelated individuals across the two sites. On the UKB dataset, we use phased autosomal haplotypes officially released in UK Biobank v3 as input to the hashing (Methods), while for All of Us, we phase a batch of 20K samples from AoU using Eagle 2 (Loh et al. 2016). We observe that independently phasing the data at each site does not affect the accuracy of SF-Relate (Supplemental Table S2).

### Ground truth preparation

To compute the ground truth of related individuals in our datasets, we follow the approach proposed in UKB documentation (Bycroft et al. 2018). We first filter the SNPs based on their implications in population structure, before computing the kinship using the KING approach (Manichaikul et al. 2010). To determine the set of SNPs to retain, we conduct a PCA on a publicly-available dataset (i.e., 1000 Genomes) using the intersection of loci with our dataset. Utilizing a reference dataset ensures that our method is not tailored to the processed dataset and effectively generalizes to other datasets. We then exclude SNPs that exhibit high PC loadings in the top three PCs, using a threshold set at the 75th percentile of these loadings. This strategy enables us to filter out SNPs with heavy loadings while retaining sufficient ancestry-agnostic autosomal SNPs for kinship inference.

Applying this approach to the UKB datasets results in selecting 90K SNPs, upon which the KING estimator predominantly identifies the same related pairs as those in UK Biobank’s relatedness release. The ground-truth relatedness degrees in our experiments are based on these KING coefficients, utilizing the recommended thresholds 2^−*d*−1.5^ for degree *d* (Manichaikul et al. 2010).

### Kinship estimation using alternative non-secure methods

Even though SF-Relate builds upon the KING estimator (Manichaikul et al. 2010), it outperforms it due to its novel approach in pre-selecting individuals likely to be related, utilizing an encoding and hashing scheme specialized to capture IBD signals. To showcase this, we compare SF-Relate and standard KING estimator alongside two advanced relatives detection tools: PC-Relate (Conomos et al. 2016) and RAFFI (Naseri et al. 2021). For PC-Relate, we rely on the hail implementation (Hail 2023) and consider only bi-allelic variants from the UK Biobank SNP panel to compute all pairwise coefficients. We set the minor allele frequency and the number of PCs to 0.1 and 10, respectively, and remove variants with missing rate higher than 5%. Additionally, we perform ldpruning on the SNP set, reducing from 600K to 300K variants using parameters *r*^2^ = 0.05 and bp window size = 500000. For kinship estimation with RAFFI, we first run the IBD-finding tool RAPID with the parameters −r3 − s1 − d5 − w3, followed by executing RAFFI (v0.1) for kinship estimation. However, we observed that the initial segment of length 10 cM on chromosome 15 is only covered by 16 base pairs in the UKB dataset, resulting in excessive candidate IBD segment pairs processed by RAPID. As the 10 cM sharing negligibly affects the overall kinship coefficient, we removed these base pairs when running RAPID on UKB.

### Phenotype simulation for the GWAS case study

To evaluate the effectiveness of SF-Relate in mitigating the confounding effects of cryptic relatedness for GWAS, we simulate a GWAS study on a subset of 100K samples from UKB using simulated phenotypes, following the methodology proposed in REGENIE (Mbatchou et al. 2021) as described next. We select a set of random variants from odd-chromosomes to serve as causal variants, reserving the even-chromosomes to assess the level of false positive associations. We exclude extremely rare SNPs (minor allele count *<* 5), before randomly selecting *P* SNPs located in odd-numbered chromosomes, for *P* ∈ {100, 1000, 10000}. These selected SNPs are designated as causal. For each causal SNP, we sample its effect size *β*_*j*_ with the constraint that the total variance (i.e., narrow-sense heritability) is *h*^2^ = 0.2. We then use a linear model with top principal component correction to simulate the phenotype *Y*_*i*_ for each individual *i* as

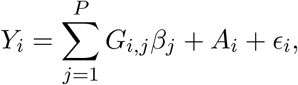

where *G*_*i,j*_ is the standardized genotype of individual *i* at SNP *j, A*_*i*_ is the first PC score of individual *i* (scaled to have variance 0.05), and *ϵ*_*i*_ is a Gaussian noise variable with variance 0.75, representing environmental effects.

### Data access

Data access application to genotypes and haplotypes from the UK Biobank can be submitted at: https://www.ukbiobank.ac.uk/. The All of Us (AoU) Controlled Tier Dataset v5 is available through the Controlled Tier of the AoU Researcher Workbench. The application to access the AoU dataset can be submitted at: https://www.researchallofus.org/register/.

## Code availability

Our open-source software, along with a demo using a public dataset, are available at: https://github.com/froelich/sf-relate. Additionally, SF-Relate can be conveniently executed through sfkit, a web server for secure collaborative genomic studies, accessible at: https://sfkit.org.

## Competing interest statements

The authors declare no competing interests.

## Acknowledgments

This work has been accepted for an oral presentation at RECOMB 2024. This work is supported by NIH R01 HG010959 (to B.B.) and NIH DP5 OD029574 and RM1 HG011558 (to H.C.) and the Broad Institute Schmidt Fellowship (to V.P.). We thank Manaswitha Edupalli, Simon Mendelsohn and Matthew Mosca for their help in processing the biobank datasets, and for integrating SF-Relate into the sfkit web server. Additionally, we would like to thank the RECOMB reviewers for their helpful suggestions.

The All of Us Research Program is supported by the National Institutes of Health, Office of the Director: Regional Medical Centers: 1 OT2 OD026549; 1 OT2 OD026554; 1 OT2 OD026557; 1 OT2 OD026556; 1 OT2 OD026550; 1 OT2 OD 026552; 1 OT2 OD026553; 1 OT2 OD026548; 1 OT2 OD026551; 1 OT2 OD026555; IAA #: AOD 16037; Federally Qualified Health Centers: HHSN 263201600085U; Data and Research Center: 5 U2C OD023196; Biobank: 1 U24 OD023121; The Participant Center: U24 OD023176; Participant Technology Systems Center: 1 U24 OD023163; Communications and Engagement: 3 OT2 OD023205; 3 OT2 OD023206; and Community Partners: 1 OT2 OD025277; 3 OT2 OD025315; 1 OT2 OD025337; 1 OT2 OD025276. In addition, the All of Us Research Program would not be possible without the partnership of its participants.

## Supplemental Materials

**Supplemental Table S1:**
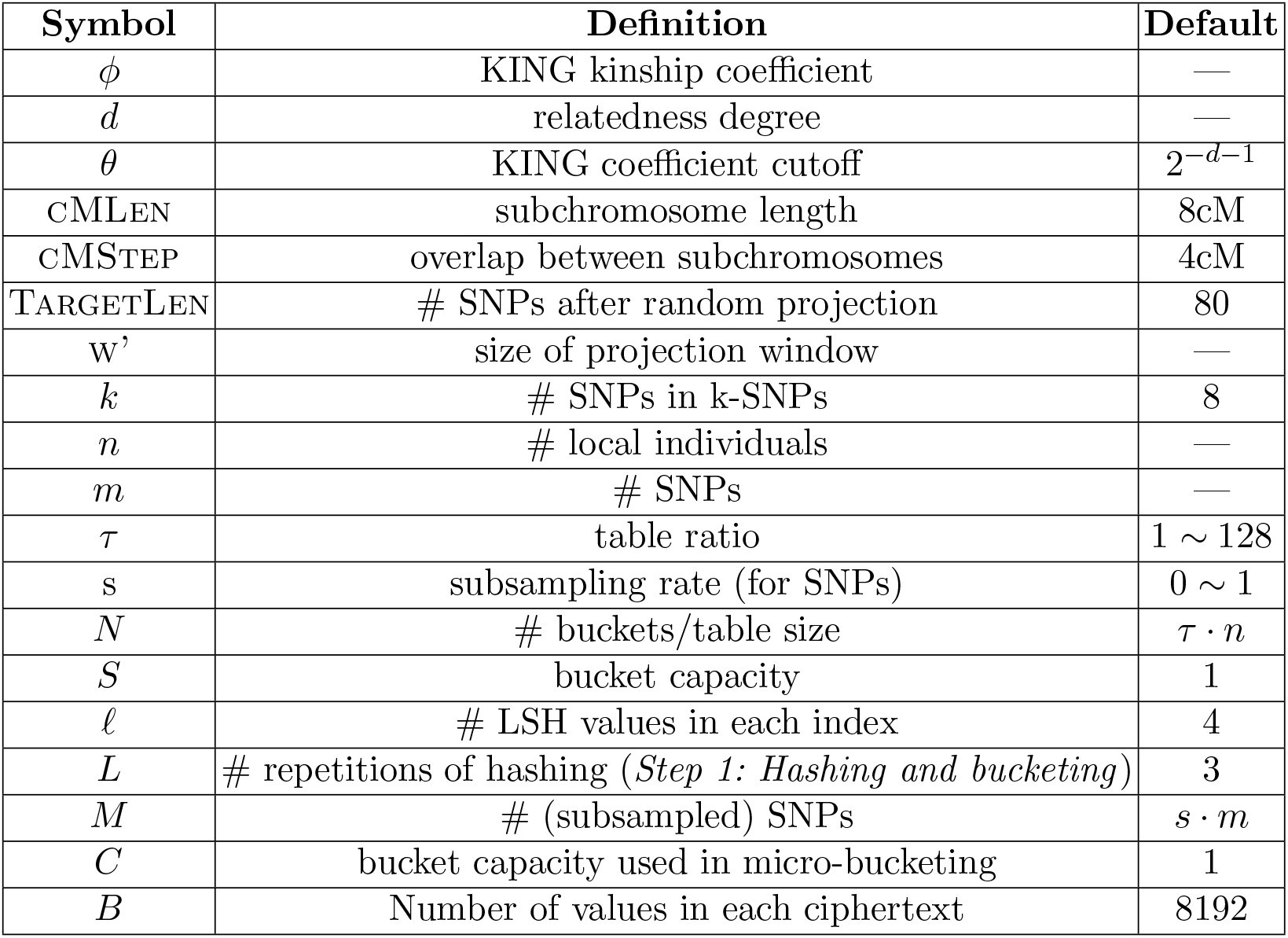
Symbols, parameters and default values. The *Default* column indicates the optimal values for the parameters across all datasets in our experiments. For the encoding pipeline (see Methods), we divide each chromosome into subchromosomes of length in equal genetic distances cMLen, with adjacent segments overlapping in a fixed distance cMStep. Empirically, 3cM to 20cM are reasonable values, with 8cM being the best for both UK Biobank and All of Us datasets. After that, we randomly project each subchromosome vector down to a vector of a fixed length TargetLen. The projection randomly selects one variant out of every window of size *w*^*′*^ SNPs, with probability proportional to their minor allele frequencies. Empirically TargetLen is set to 80 to ensure enough SNPs are chosen. To make each k-SNP correspond to roughly the same genetic distance, *w*^*′*^ is chosen differently for each subchromosome, and it is computed as the ratio between the actual number of SNPs in the subchromosome and TargetLen. The parameter *k* in k-SNP can be between 5 and 30, and 8 is the best value for all datasets. The repetition parameter *L* specifies the number of times parties should repeat *Step 1: Hashing and bucketing* (Methods). To ensure resulting merged table is highly utilized (*>* 99.9% non-dummies), *L* can be chosen on the fly independently by each party.

| Symbol | Definition | Default |
| --- | --- | --- |
| $\phi$ | KING kinship coefficient | — |
| $d$ | relatedness degree | — |
| $\theta$ | KING coefficient cutoff | $2^{-d-1}$ |
| CMLen | subchromosome length | 8cM |
| CMSTEP | overlap between subchromosomes | 4cM |
| TARGETLEN | # SNPs after random projection | 80 |
| $w'$ | size of projection window | — |
| $k$ | # SNPs in k-SNPs | 8 |
| $n$ | # local individuals | — |
| $m$ | # SNPs | — |
| $\tau$ | table ratio | $1 \sim 128$ |
| $s$ | subsampling rate (for SNPs) | $0 \sim 1$ |
| $N$ | # buckets/table size | $\tau \cdot n$ |
| $S$ | bucket capacity | 1 |
| $\ell$ | # LSH values in each index | 4 |
| $L$ | # repetitions of hashing (*Step 1: Hashing and bucketing*) | 3 |
| $M$ | # (subsampled) SNPs | $s \cdot m$ |
| $C$ | bucket capacity used in micro-bucketing | 1 |
| $B$ | Number of values in each ciphertext | 8192 |

**Supplemental Table S2:** SF-Relate effectively detects relatives across diverse datasets. We showcase the robustness of SF-Relate’s in two scenarios. (1) *Geographically Distributed*: a collaborative setting involving six centers collecting patient data within their respective geographic region (see Supplementary Table 2). The dataset comprises 100K individuals of white British ethnicity sourced from the UK Biobank. (2) *Single African Ancestry*: a collaborative setting with two parties each holding 10K samples of African ancestry from All of Us. Additionally, we confirm that using locally-phased data does not affect SF-Relate’s ability to identify relatives. We consider two examples, each involving 20K patients split between two parties and sourced either from the UK Biobank (*Locally-Phased UKB*) or All of Us (*Locally-Phased AoU*). We test two prominent phasing tools that we use independently on each site. We apply SHAPEIT 5 (`phase_common`) on UKB and Eagle v2.4.1 with the option `--maxMissingPerSnp` 0.5 on AoU. Across all settings, SF-Relate’s recall remains consistently high, all above 87%. We observe a possible minor reduction in the third-degree recall on locally phased AoU, likely due to phasing errors associated with the small dataset size. We expect local phasing to be more accurate in a larger cohort or when large public reference panels are used.

| Dataset | Recall (%, counts) |  |  |  |  |
| --- | --- | --- | --- | --- | --- |
|  | Relatedness degree |  |  |  | Overall |
|  | 0th | 1st | 2nd | 3rd |  |
| Geographically Distributed | 100.0%<br>22/22 | 100.0%<br>3370/3370 | 100.0%<br>1483/1483 | 99.5%<br>10591/10640 | 99.7%<br>15515/15466 |
| Single African Ancestry | 100.0%<br>104/104 | 100.0%<br>1278/1278 | 100.0%<br>850/850 | 98.8%<br>530/542 | 99.6%<br>3262/3274 |
| Locally-Phased UKB | 100.0%<br>16/16 | 99.5%<br>3999/4017 | 98.2%<br>1598/1626 | 93.2%<br>3604/3863 | 96.7%<br>9217/9522 |
| Locally-Phased AoU | 100.0%<br>14/14 | 100.0%<br>209/209 | 97.8%<br>91/93 | 87.0%<br>134/154 | 95.3%<br>434/468 |

**Supplemental Table S3:**
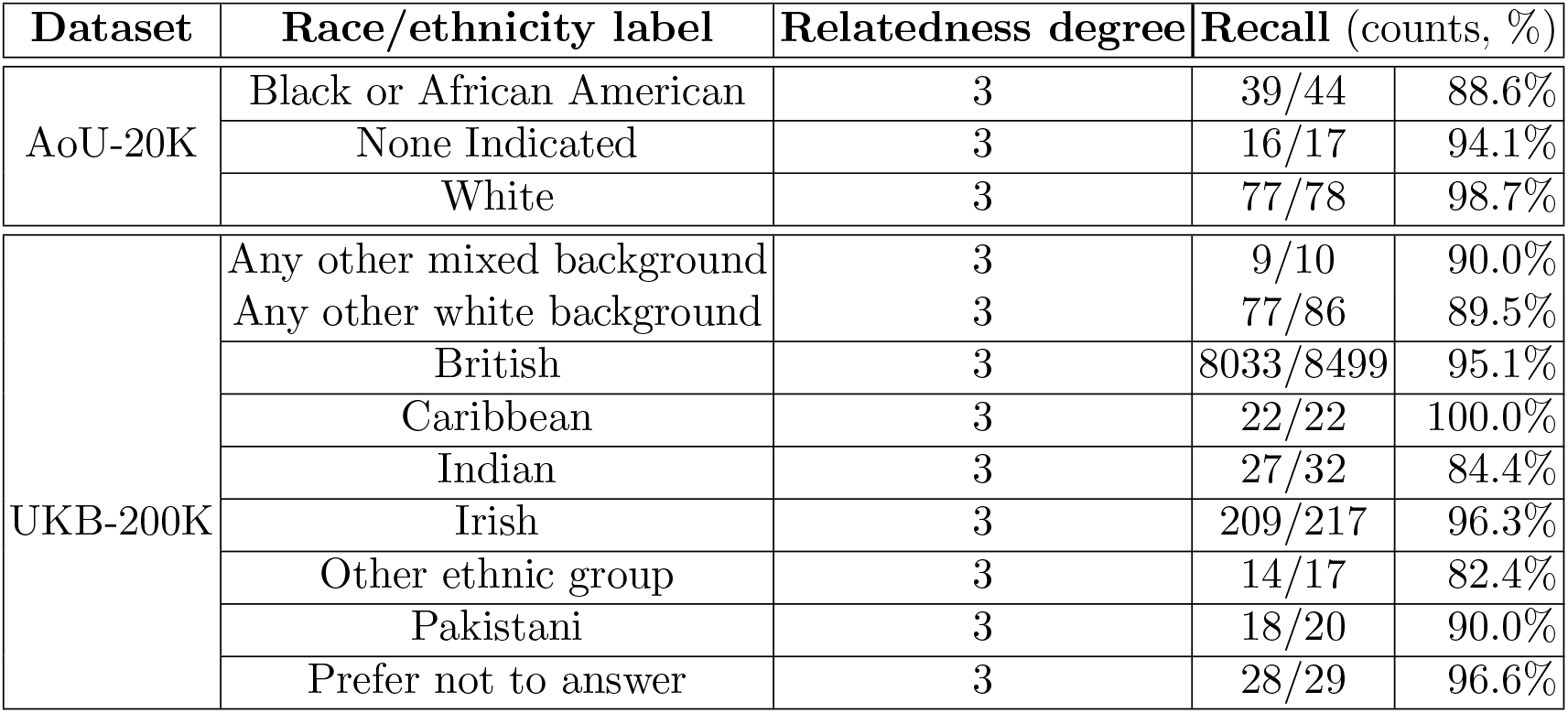
SF-Relate’s recall remains high across different subpopulations. We apply SF-Relate to the multi-ethnicity datasets with 20K individuals from AoU and 200K individuals from UKB, each split between two parties. The recall for the 0th, 1st and 2nd degrees is nearly 100% for all subpopulations in both datasets, and thus excluded from the table. We separately calculate and report the third-degree recall for each subpopulation with at least 20 third-degree samples.

**Supplemental Table S4:** SF-Relate achieves high recall across various parameter settings. On UKB-200K, we select a combination of reasonable parameters and compute the third-degree recall achieved by SF-Relate. The overlap between chromosomes cMSTEP is set to equal half of the subchromosome length cMLEN (see Supplemental Table S1). We vary the values of *k*, the number of SNPs in *k*-SNPs and *ℓ* the number of LSH values in each index, and maintain the remaining parameters at their default values provided in Supplemental Table S1. SF-Relate consistently achieves near-perfect recall (*>* 95%), except on the lower end of the parameters, when there is not enough entropy among the subchromosomes for the LSH to separate samples into buckets.

|  |  |  |  |  |  |  |  |  |  |
| --- | --- | --- | --- | --- | --- | --- | --- | --- | --- |
| cMLEN | 5.0 |  |  |  |  |  |  |  |  |
| $k$ | 6.0 | | | 8.0 | | | 10.0 | | |
| $\ell$ | 3.0 | 4.0 | 5.0 | 3.0 | 4.0 | 5.0 | 3.0 | 4.0 | 5.0 |
| **Recall (%)** | **84.2** | **96.6** | **97.5** | **96.1** | **97.5** | **98.3** | **97.6** | **98.0** | **98.3** |
| cMLEN | 8.0 |  |  |  |  |  |  |  |  |
| $k$ | 6.0 | | | 8.0 | | | 10.0 | | |
| $\ell$ | 3.0 | 4.0 | 5.0 | 3.0 | 4.0 | 5.0 | 3.0 | 4.0 | 5.0 |
| **Recall (%)** | **82.7** | **96.0** | **97.3** | **96.0** | **97.9** | **98.1** | **97.7** | **98.0** | **98.5** |
| cMLEN | 11.0 |  |  |  |  |  |  |  |  |
| $k$ | 6.0 | | | 8.0 | | | 10.0 | | |
| $\ell$ | 3.0 | 4.0 | 5.0 | 3.0 | 4.0 | 5.0 | 3.0 | 4.0 | 5.0 |
| **Recall (%)** | **77.3** | **95.7** | **97.2** | **96.0** | **97.3** | **97.5** | **97.4** | **97.5** | **97.9** |

**Supplemental Table S5:** SF-Relate outperforms KING and achieves comparable results to more advanced methods, PC-Relate and RAFFI. On a subset of 20K samples from UKB-200K, we compute the maximum relatedness degree for each individual using the different methods, and report the number of individuals falling into each classification type (when this count exceeds 20). The abbreviation U represents *Unrelated*. **<u>Underlined columns</u>** correspond to the set of individuals with identical classifications across all methods. **Bold-faced columns** denote substantial sets containing at least 100 individuals. Most methods have consistent results in the largest categories, except in column I. Whereas SF-Relate and KING agree on most columns, SF-Relate outperforms KING by aligning with the more accurate methods in column I, effectively excluding the thousands of likely-spurious 4th-degree pairs identified by KING.

| Degree combinations → | <u>A</u> | <u>B</u> | <u>C</u> | D | <u>E</u> | F | G | H | **I** | J | <u>K</u> |
| --- | --- | --- | --- | --- | --- | --- | --- | --- | --- | --- | --- |
| SF-Relate | <u>1st</u> | <u>2nd</u> | <u>3rd</u> | 3rd | <u>4th</u> | 4th | U | U | **U** | U | <u>U</u> |
| KING | <u>1st</u> | <u>2nd</u> | <u>3rd</u> | 3rd | <u>4th</u> | 4th | 3rd | 4th | **4th** | U | <u>U</u> |
| RAFFI | <u>1st</u> | <u>2nd</u> | <u>3rd</u> | 4th | <u>4th</u> | U | 3rd | 4th | **U** | U | <u>U</u> |
| PC-Relate | <u>1st</u> | <u>2nd</u> | <u>3rd</u> | 3rd | <u>4th</u> | U | 3rd | 4th | **U** | 4th | <u>U</u> |
| Number of individuals | **508** | **173** | **863** | 48 | **93** | 23 | 22 | 23 | **8410** | 23 | **9716** |

**Supplemental Table S6:** Assignment of UK Biobank assessment centers to geographic areas. To simulate a federated study, we use a subset of 100K white British individuals from UK Biobank dataset. We organized the 22 data collection centers (Data-Field 54) into six study groups according to their geographic locations within the UK.

| UK Biobank Assessment Center | Geographic Area | Number of Samples |
| --- | --- | --- |
| Edinburgh<br>Glasgow | Scotland | 7051 |
| Middlesbrough<br>Newcastle | Northeast England | 14185 |
| Liverpool<br>Bury<br>Stockport<br>Manchester<br>Leeds | Northwest England | 25096 |
| Birmingham<br>Stoke<br>Sheffield<br>Nottingham<br>Wrexham | Southeast England | 23362 |
| Barts<br>Hounslow<br>Croydon<br>Reading<br>Oxford | Central England | 16836 |
| Swansea<br>Cardiff<br>Bristol | Wales | 13470 |
| **Total Number of Samples** |  | 100,000 |

**Supplemental Figure S1:**
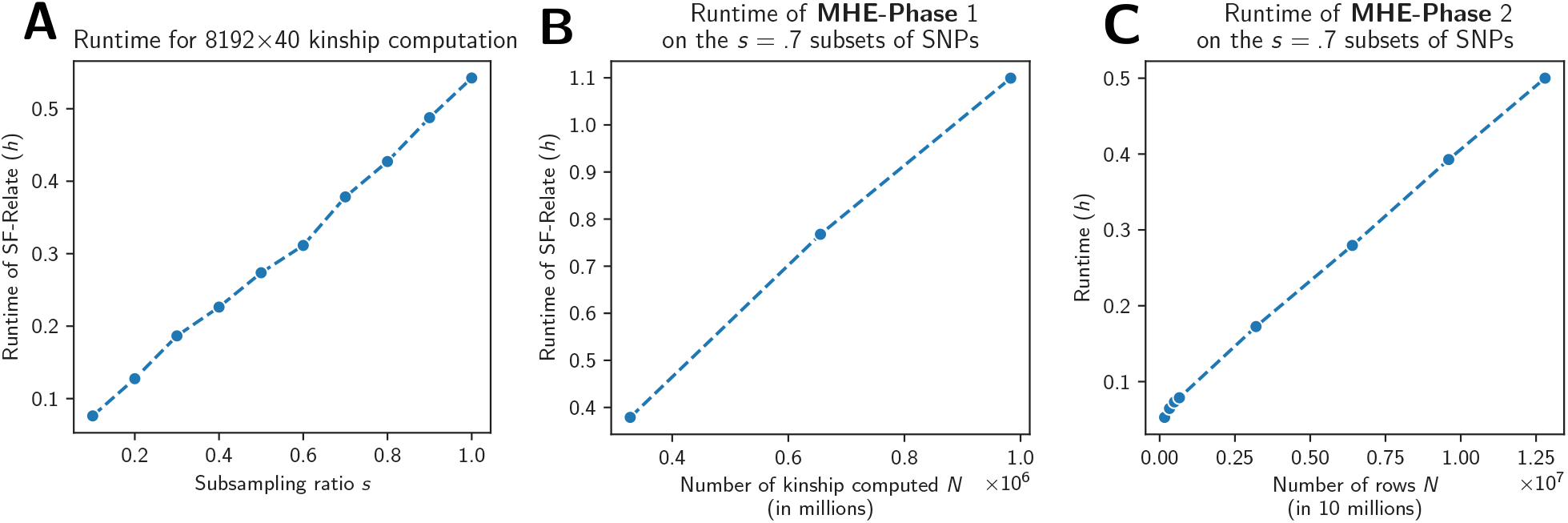
SF-Relate’s runtime scales linearly in database dimension in practice. In **A**, we perform a fixed number of kinship computations *N* while varying the number of SNPs by varying the subsampling rate *s*. In **B** and **C**, we increase the number of kinship computations *N* while keeping the subsampling rate *s* at .7. We report the runtime of the MHE-Phase 1 and MHE-Phase 2 in (B) and (C) separately.

**Supplemental Figure S2:**
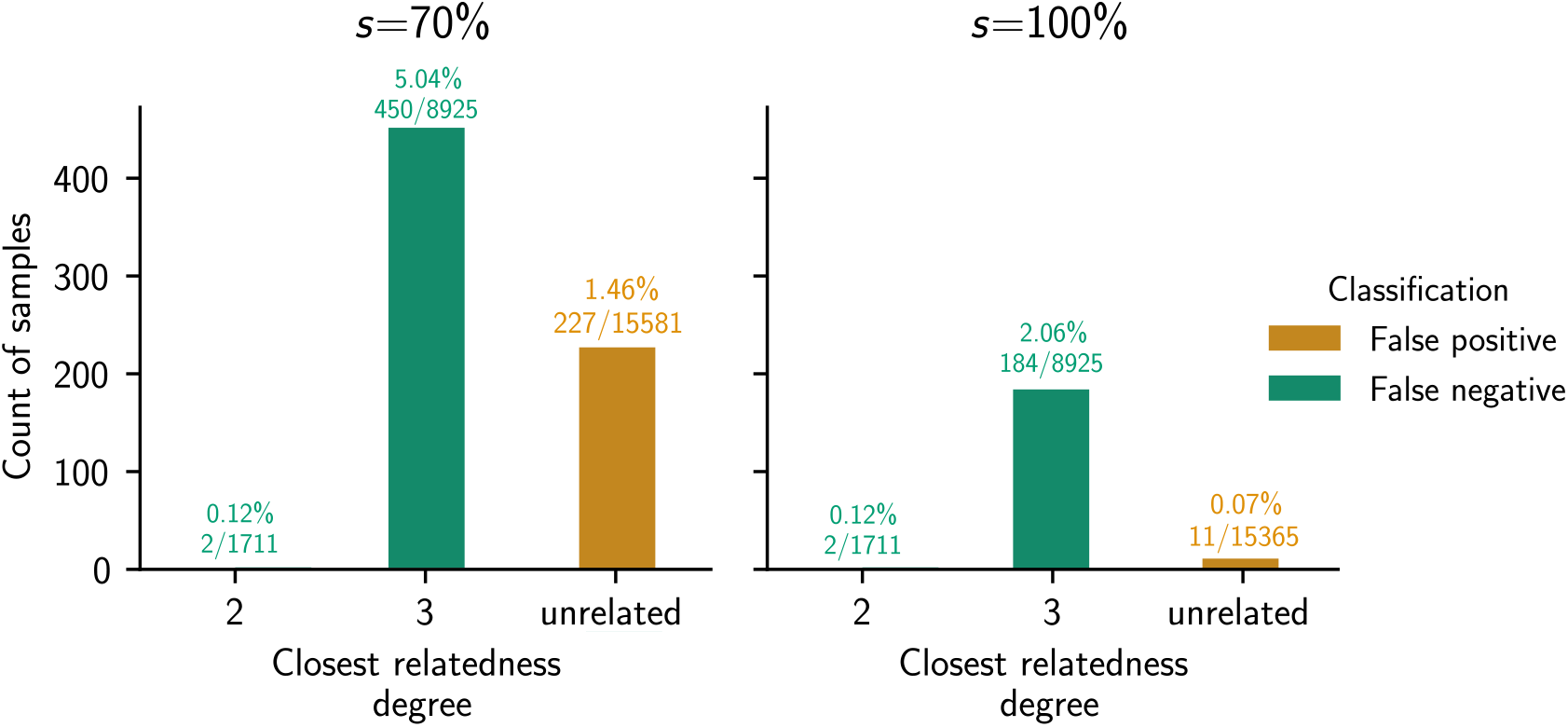
Increasing sketching ratio *s* allows almost perfect precision. We perform SF-Relate on UKB-200K under the alternative sketching (subsampling) ratio *s* = 1, and shows counts of individuals based on the their closest relations and SF-Relate’s detection results. Using the full set of SNPs in SF-Relate allows significant reductions in false positives and false negatives due to the sketching noise, with a moderate increase of runtime from 14.5 to 21 hours.

**Supplemental Figure S3:**
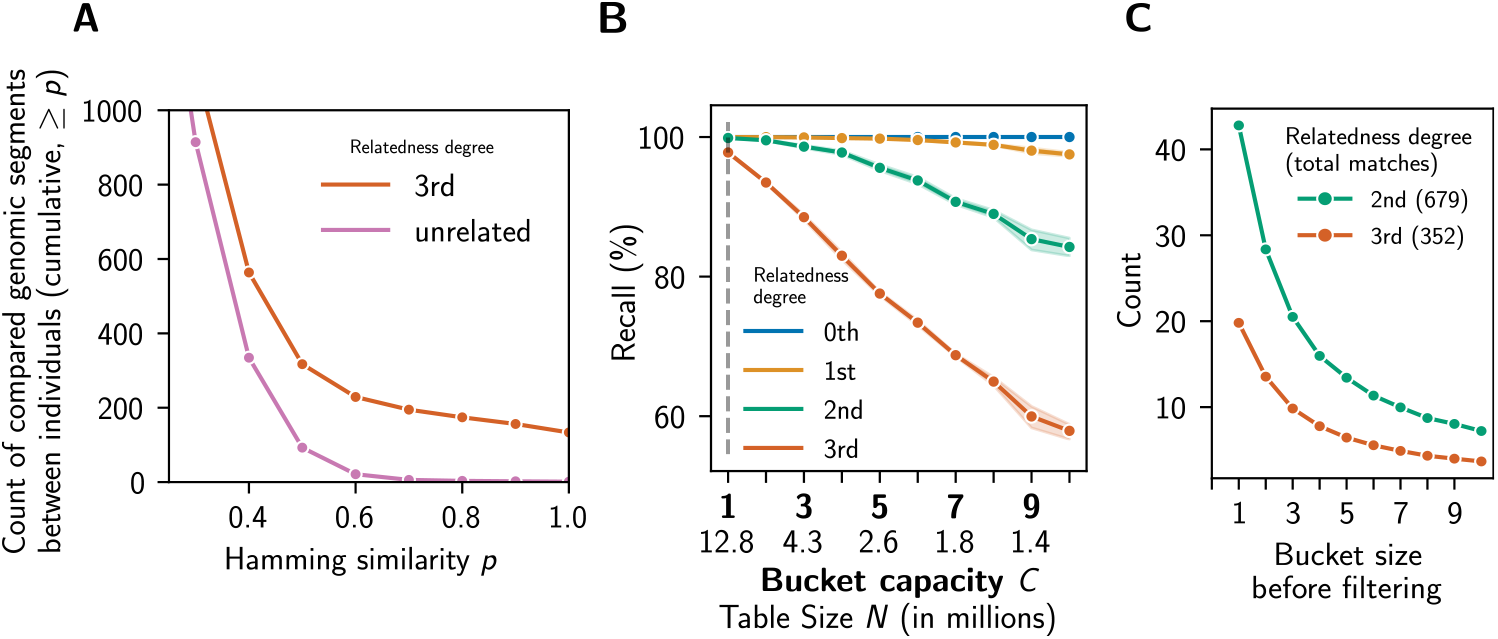
SF-Relate’s hashing and micro-bucketing strategies effectively assign close relatives to the same bucket. (**A**) Hamming LSH with SF-Relate’s k-SNP encoding scheme enables separation of pairs of genomic segments with high Hamming similarity (likely IBD) between close relatives from those pairs between unrelated individuals. (**B**) Setting the bucket capacity *C* = 1 achieves the highest recall in relative detection compared to larger values of *C*. For comparison, we adjust the table size *N* accordingly to keep the number of comparisons *NC*^2^ = 1.28 million constant. The recall for each relatedness degree is the fraction of relative pairs of that degree that are assigned to the same bucket. (**C**) Close relatives (of 2nd and 3rd degrees) who are assigned to the same bucket are most often found in size-1 buckets before trimming. All results are based on UKB-200K.

**Supplemental Figure S4:**
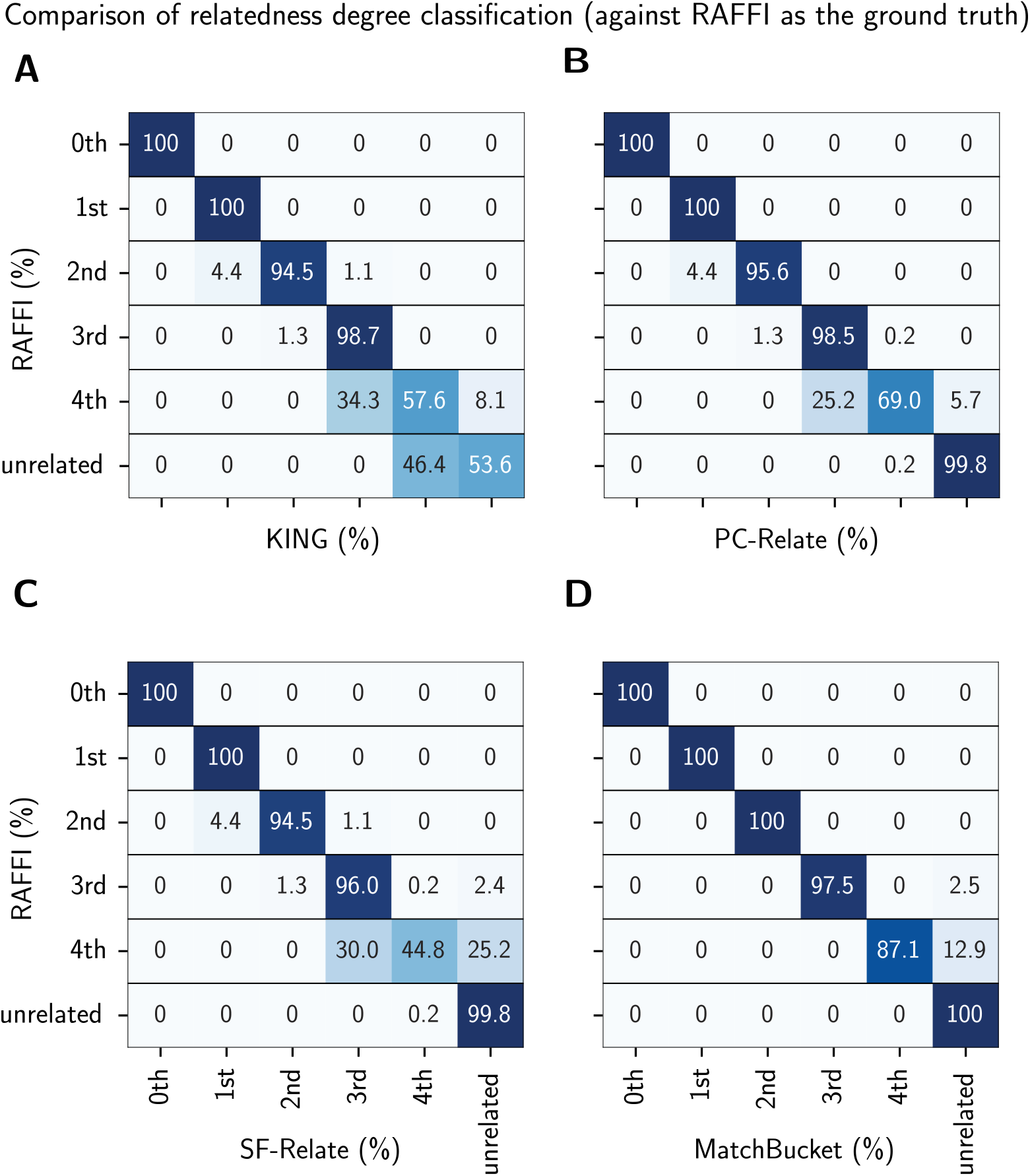
SF-Relate and PC-Relate exclude spurious 4th-degree relatives detected by KING, when compared to RAFFI as the ground-truth. On a subset with 20K samples from UKB-200K, we present the confusion matrices assessing the relatedness classification accuracy of KING **(A)**, PC-Relate **(B)** and SF-Relate **(C)**, comparing them with the output of RAFFI as the ground-truth. *MatchBucket* **(D)** denotes the (non-private) hybrid approach where RAFFI is performed in plaintext on pairs in SF-Relate’s corresponding buckets and serves as a reference. Both SF-Relate and PC-Relate label RAFFI-unrelated individuals as unrelated, unlike KING, which labels them as 4th-degree relatives. This suggests that both methods avoid the spurious relationships identified by KING.

**Supplemental Figure S5:**
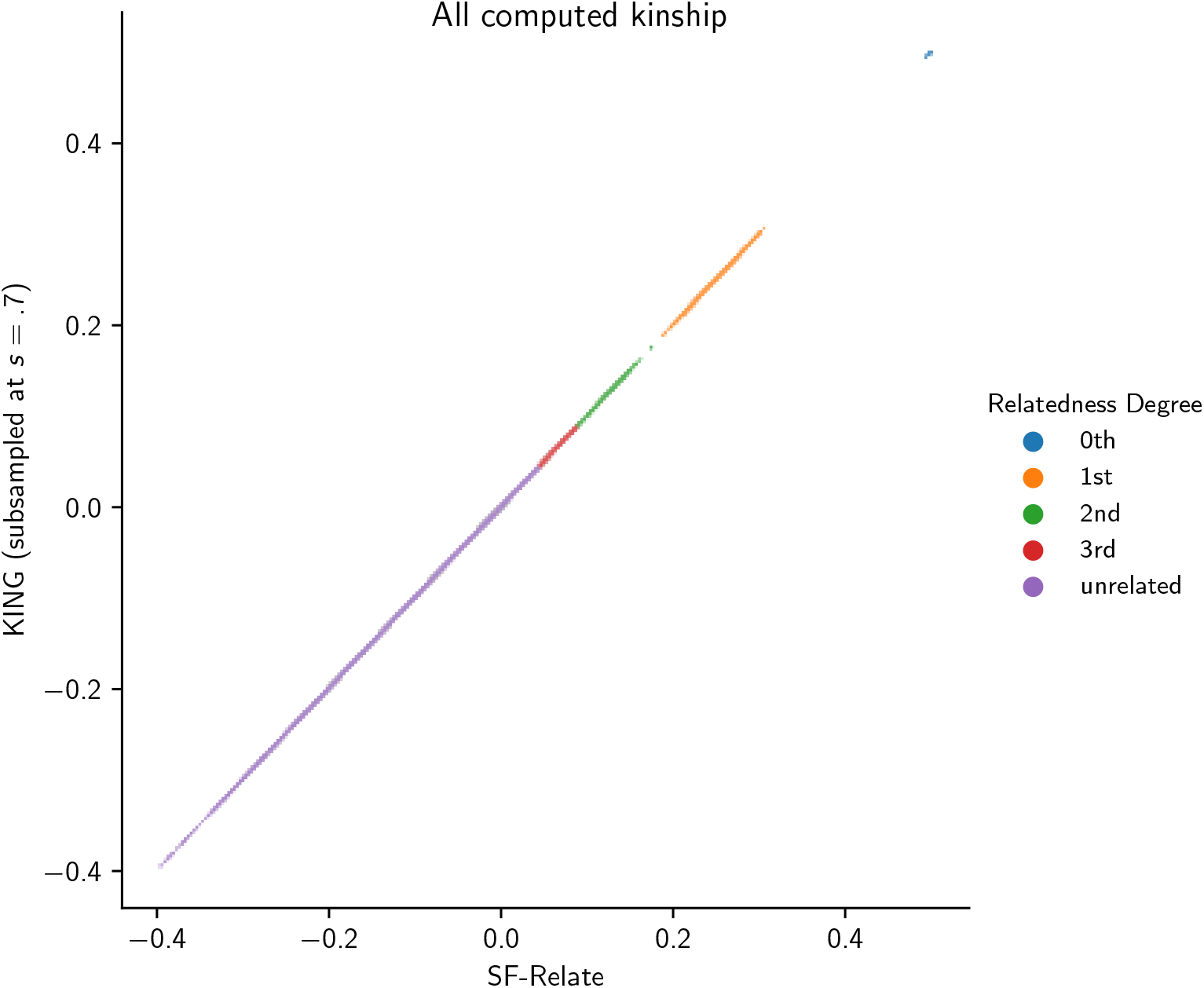
SF-Relate’s alternative output mode accurately reports all computed kinship coefficients. On a 10% random subset of kinship coefficients that SF-Relate computed on UKB-200K, we evaluate the accuracy of the alternative setting of SF-Relate where the full list of kinship coefficients is revealed. As shown in the plot, SF-Relate’s output accurately matches with KING, indicating that the MHE noise is negligible with respect to the kinship coefficients being computed on the subsampled set of SNPs.

**Supplemental Figure S6:**
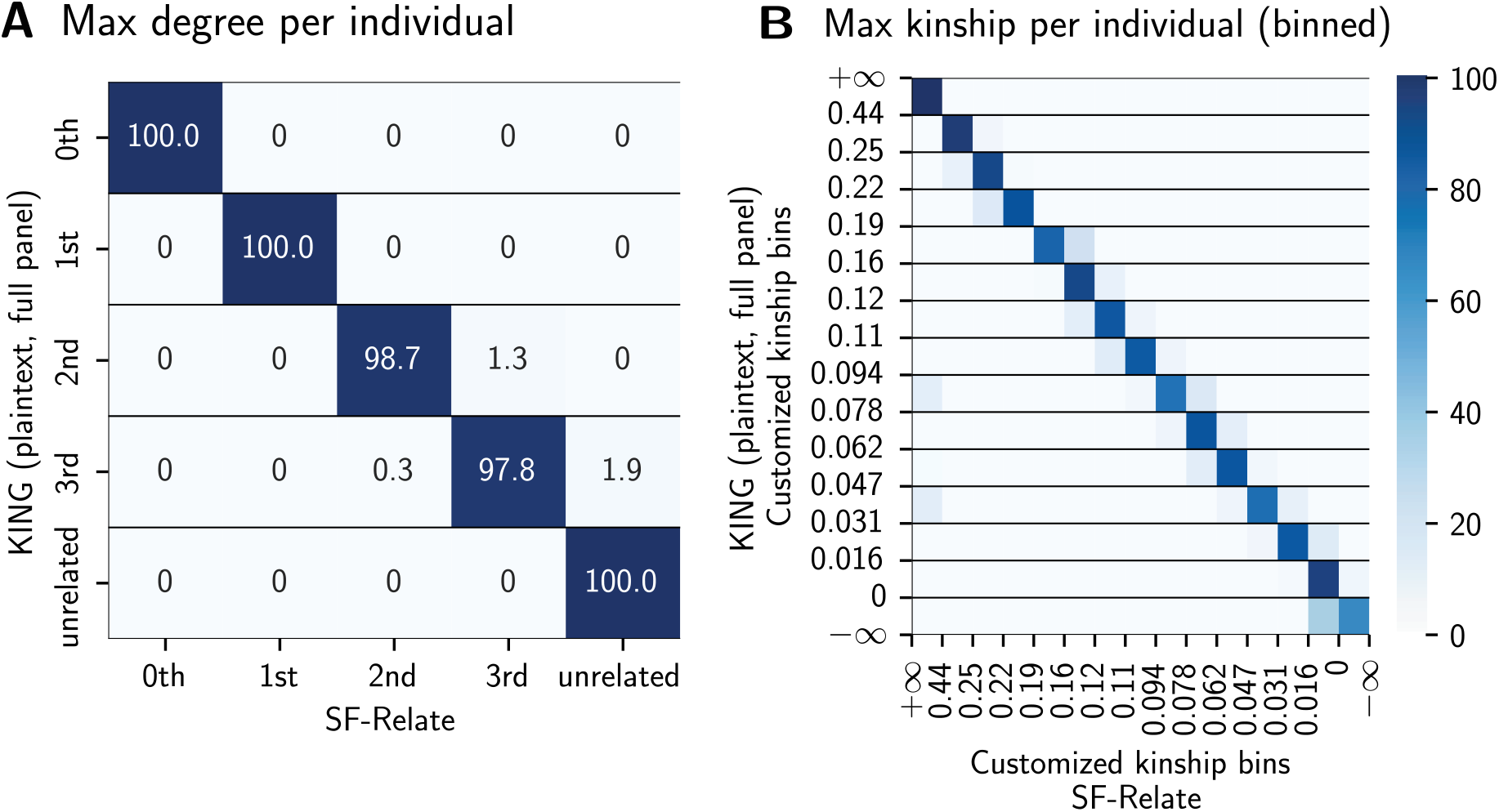
SF-Relate’s alternative output modes compute accurate individuals-level statistics with customizable thresholds. On a subset of 10% kinship coefficients computed in UKB-200K’s hash tables, we evaluate SF-Relate’s alternative output modes that support the comparison of the computed kinship coefficients with a sequence of thresholds. Numbers in the cells or their colors indicate the corresponding recall rates. In **(A)**, we choose the thresholds to be the recommended kinship cutoff by KING in Manichaikul et al. (2010). In **(B)**, the thresholds defining the refined bins are marked on the axes. The predictions perfectly align with the ground-truth KING predictions computed on the full SNP panel on more than 99.9% and 85% individuals in **(A)** and **(B)**, respectively. More than 99.9% of the predictions in **(B)** are accurate, with deviations at most shifted by a single bin. This result indicates that both modes produce reliable assessment of the maximum kinship level, and highlight the utility of SF-Relate in various workflows.

**Supplemental Figure S7:**
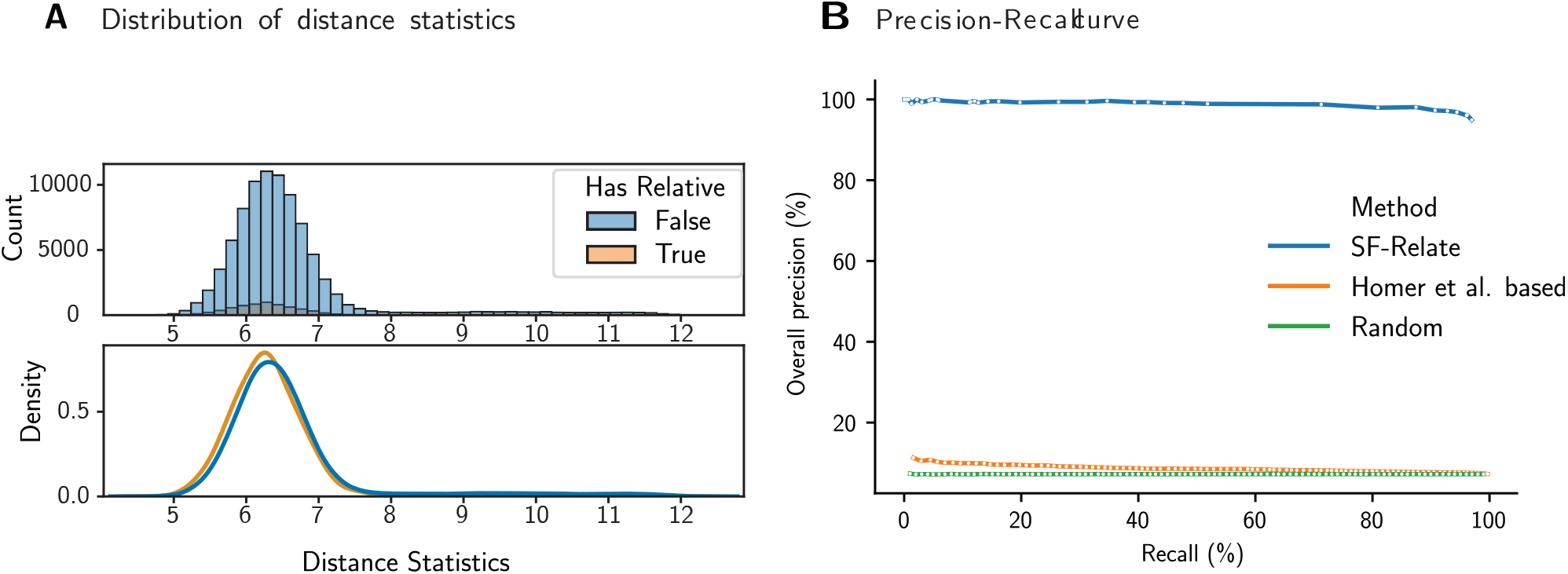
Relative detection based on Homer et al.’s attack results in near-random performance. Homer et al.’s attack (2008) predicts whether an individual contributes to a genetic dataset by statistically testing whether the individual’s genotype count vector is closer to the allele frequencies in the dataset than some alternative reference frequencies. The same attack can potentially reveal the presence of a relative in a dataset due to shared genetic sequences. We evaluate the efficacy of this approach on the UKB-200K dataset split among two parties. In (**A**), we compute (1) the *L*^1^ distance of every sample’s genotype count vectors to the mean genotype count vector from the local dataset (excluding relatives), representing the background statistic, and (2) the distance between the sample genotype count vector and the mean count vector in the other party’s dataset, representing the target statistic. We then subtract the two distances to see which dataset the sample is closer to. The figure shows that this estimator does not effectively separate samples that have a relative in the other dataset from samples that do not. In (**B**), we show the precision-recall curves for various methods for detecting 3rd-degree or closer relatives in UKB-200K. For Homer et al., we vary the distance threshold used as a cut-off to determine whether a sample is closer to the target dataset. For SF-Relate, we plot the curve by varying the table size parameter (*τ* in Methods). Homer et al.’s approach obtains precision comparable to the level of random guessing, whereas SF-Relate achieves near-perfect precision.

**Supplemental Figure S8:**
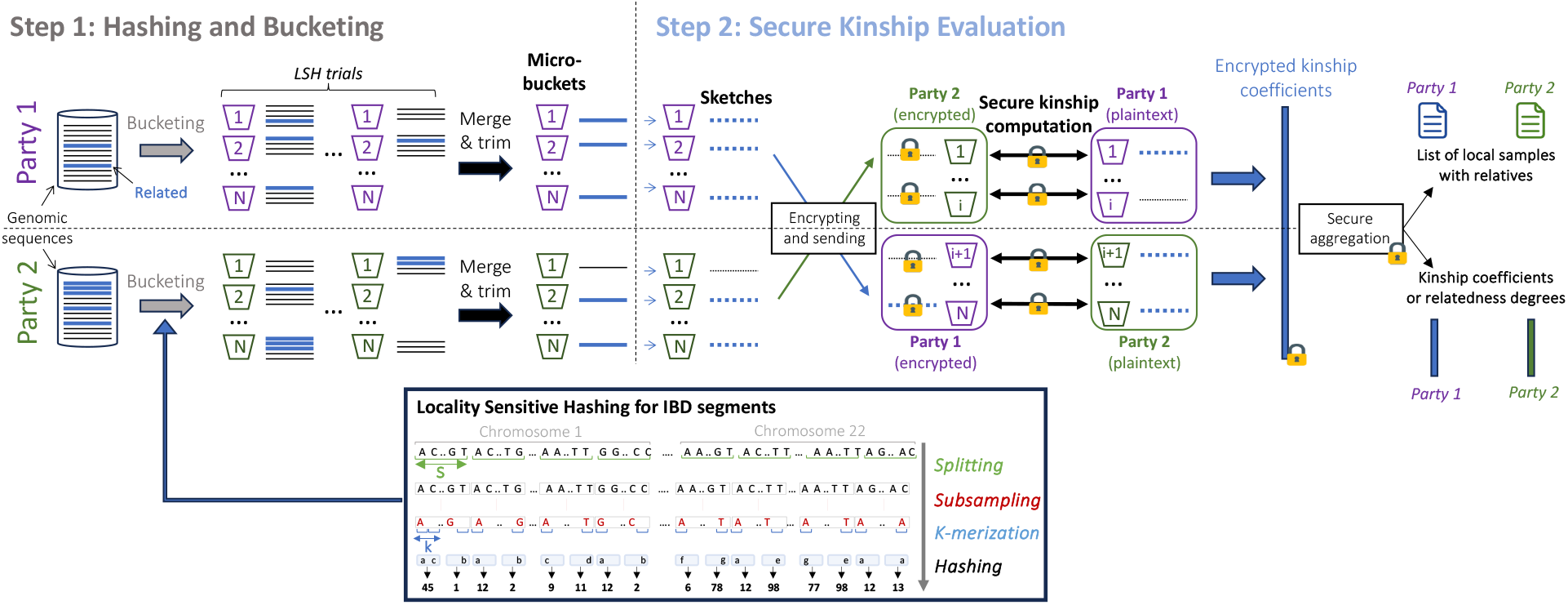
SF-Relate’s workflow. In Step 1, the parties perform multiple trials of LSH to bucket samples before merging and trimming to obtain buckets of size 1 (micro-buckets). In Step 2, each sample is sketched and securely compared against the other party’s sample in the same bucket to evaluate kinship (MHE-Phase 1). Finally, parties securely aggregate the results to obtain per-sample output (MHE-Phase 2).

**Supplemental Figure S9:**
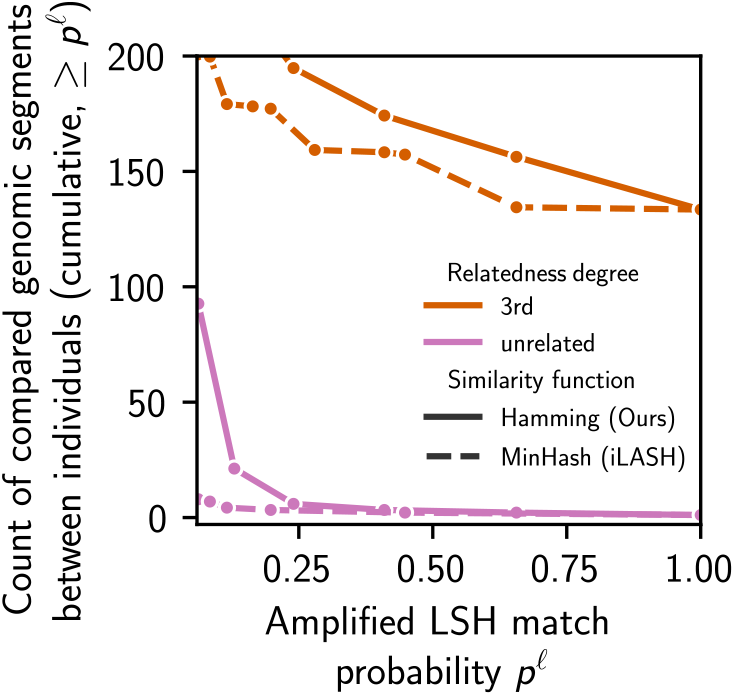
The LSH for Hamming similarity retains more high-probability pairs (candidate IBD segments) than LSH for Jaccard similarity (Min-Hash) On the UKB-200K dataset, we count the number of subchromosome pairs between individuals based on the probability that the pairs would produce the same LSH index with an LSH amplification with *ℓ*= 4. A higher probability between segments indicate higher similarity and thus more likely to be in IBD. The raw similarity score 0 ≤ *p* ≤ 1 between vectors is transformed to the amplified LSH match probability *p*^l^ by the amplification described in *Step 1: Hashing and bucketing* (Methods). Each curve shows the count of segments with probability higher than *p*^*ℓ*^ averaged over pairs of related samples in the respective relatedness degree classes. The plot shows that Hamming LSH identifies more segments with high matching probability after the LSH amplification, suggesting its ability to detect more IBD segments.

**Supplemental Figure S10:**
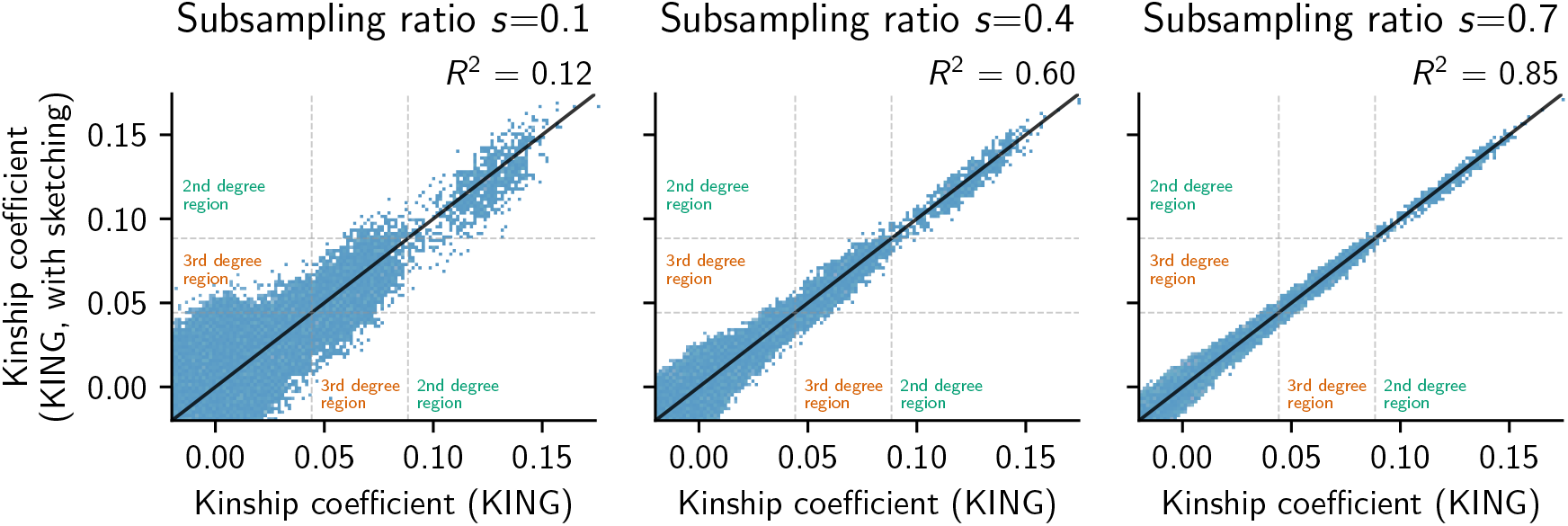
KING can be accurately estimated on subsampled subsets of SNPs. A subsampling ratio at *s* = .7 achieves reliable relatedness degree classification, where accurate kinship is not necessary. We apply SF-Relate on UKB-200K and compute the kinship coefficients on the list of micro-bucket pairs, under subsets of SNPs with different subsampling ratios. The *x* axis shows the kinship on the full set of SNPs. Points near the degree boundaries have a higher chance of classification error. The scatterplots show that sketching introduces noise to the kinship, but reliable relatedness degree classification is possible when points are not too close to the thresholds.

**Supplemental Note S1 Secure MHE protocols for secure kinship evaluation**

We detail here SF-Relate protocols for computing and detecting kinship coefficients that are above a predefined threshold (MHE-Phase 1) and for aggregating these results per individual (MHE-Phase 2).

We denote a matrix ***A*** with *N* rows and *M* columns as ***A***^*N×M*^ and a vector ***x*** with *B* elements as ***r***^*B×*1^. We use 0-based indexing, i.e. columns and rows are numbered starting from 0. **x**[*a*: *b*] denotes the subvector from row *a* to *b* (including *a* but excluding *b*). We omit *a* when it is 0, and omit *b* if it equals *N* − 1. Similarly, for matrices, we use **A**[*a*: *b, c*: *d*] to denote the submatrix specified by the ranges. For a vector ***x***^*N×*1^ = (*x*_0_, …, *x*_*N*−1_), we use ***x***^2^ to denote the vector 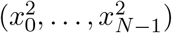, i.e. the one where elements are squared. The notation Sign(*E*) denotes the indicator of the event *E*. When applied to a vector of events as in Sign(***x***^*n×*1^ == 1), it corresponds to the vector of indicators. In other words, Sign(***x*** == 1) is equivalent to (Sign(***x***[*i*] == 1), …, Sign(***x***[*n* − 1] == 1)). Finally, **0**^*B×*1^ and **1**^*B×*1^ denotes the vector (0, 0, …, 0)^*B×*1^ and (1, 1, …, 1)^*B×*1^, respectively.

Every ciphertext under the CKKS encryption (Cheon et al. 2017) encrypts a vector with length *B*, the CKKS block length, which is typically a power of 2 like 8192. We denote the encryption of a vector ***x*** as 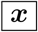. We simplify the function interfaces of the CKKS implementation in Lattigo (2022) as follows. Cryptographic keys are omitted in the function calls.

– Enc(**x**) takes in a plaintext vector **x** and returns an encrypted ciphertext 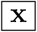.
– CollaborativeDecrypt 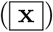 takes in an encrypted vector 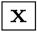 and returns the plaintext result **x**. Note that all parties would need to collaborate for this operation.
– Homomorphic Single-Instruction Multiple-Data (SIMD) operations, including coordinate-wise addition, subtraction and multiplication between any combination of ciphertexts encrypting vectors and plaintext vectors are denoted by +,−, ·, respectively.
– The Rotate 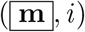 operation that that receives as input **m**^*B×*1^ = (*m*_0_, … *m*_*B*−1_) outputs a ciphertext encrypting (*m*_*i*_, *m*_*i*+1_, …, *m*_*B*_, *m*_0_, *m*_1_, …, *m*_*i*−1_).

We also implemented the following helper functions:

– Sign 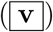 makes use of the Chebyshev polynomial interpolation and Newton’s method to approximately compute the sign function, using combinations of the homomorphic SIMD operations +, − and ·. It returns a ciphertext 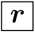 such that if one decrypts it, the result equals the Boolean-valued indicator for Sign(**v**).
– Extract 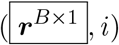 returns a ciphertext encrypting the vector (0, …, 0, ***r***[*i*], 0, …, 0)^*B×*1^. We implement this by homomorphically multiplying the *i*-th basis vector with 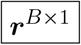.
– InnerSum 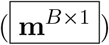 transforms the vector **m** = (**m**[0], …, **m**[*B* − 1]) into the vector **s** = (*s, s*, …, *s*)^*B×*1^ where 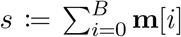. Adding a length-*B* vector with its powers-of-2 rotated copy (see Rotate(***m***, *i*) above), for all the powers of 2 at most *B*, namely 2^1^, 2^2^, …, *B* achieves this. Hence, we implement it efficiently by iterating over 0, …, log *B* rotating ***m*** accordingly, and then homomorphically add the results together.

We display here the two protocols used in Methods, namely MHE-Phase 1 and MHE-Phase 2, for secure kinship evaluation between two parties.

### MHE-Phase 1

Distributed relative detection

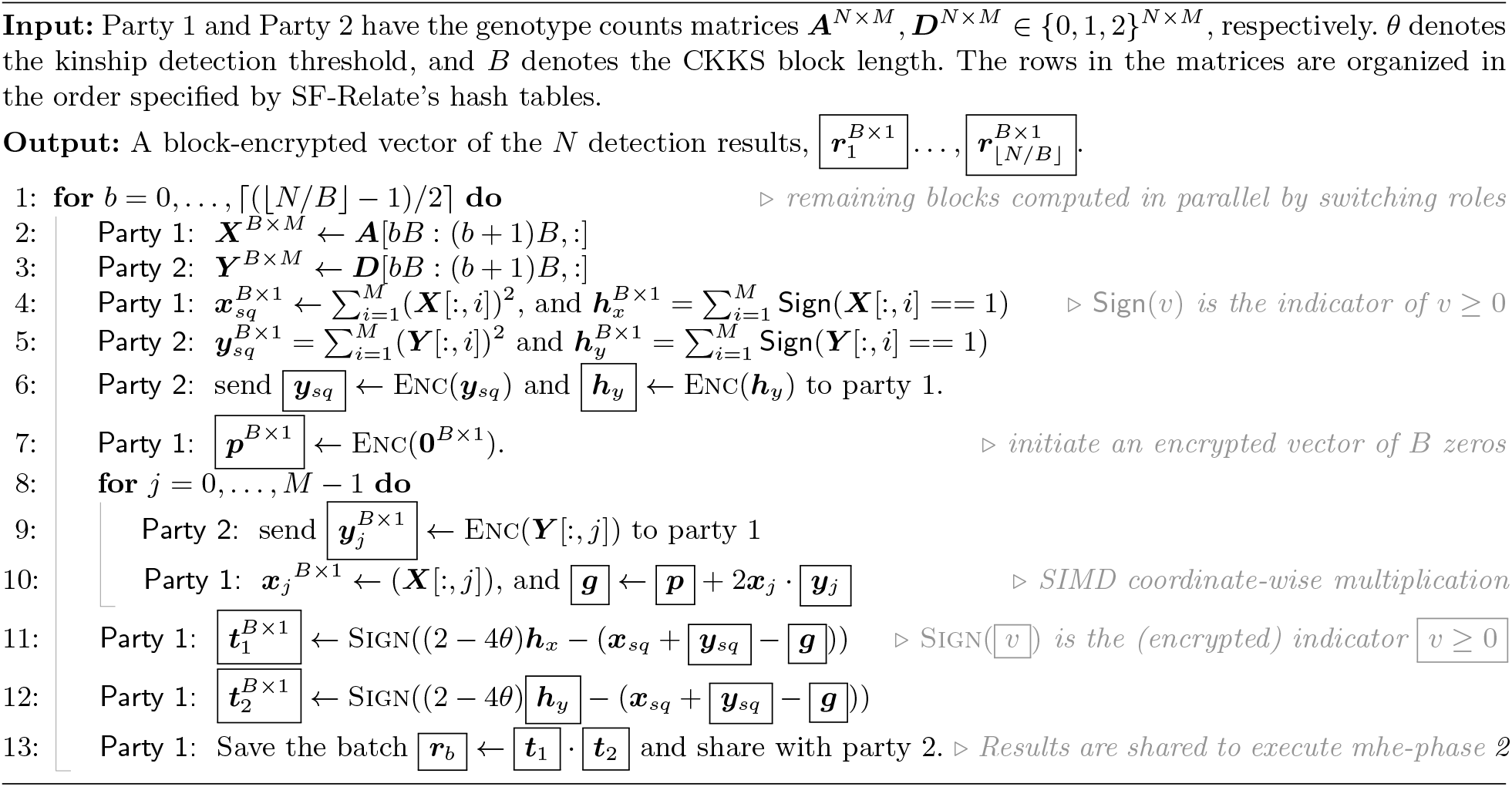

### MHE-Phase 2

Accumulation to per-sample output

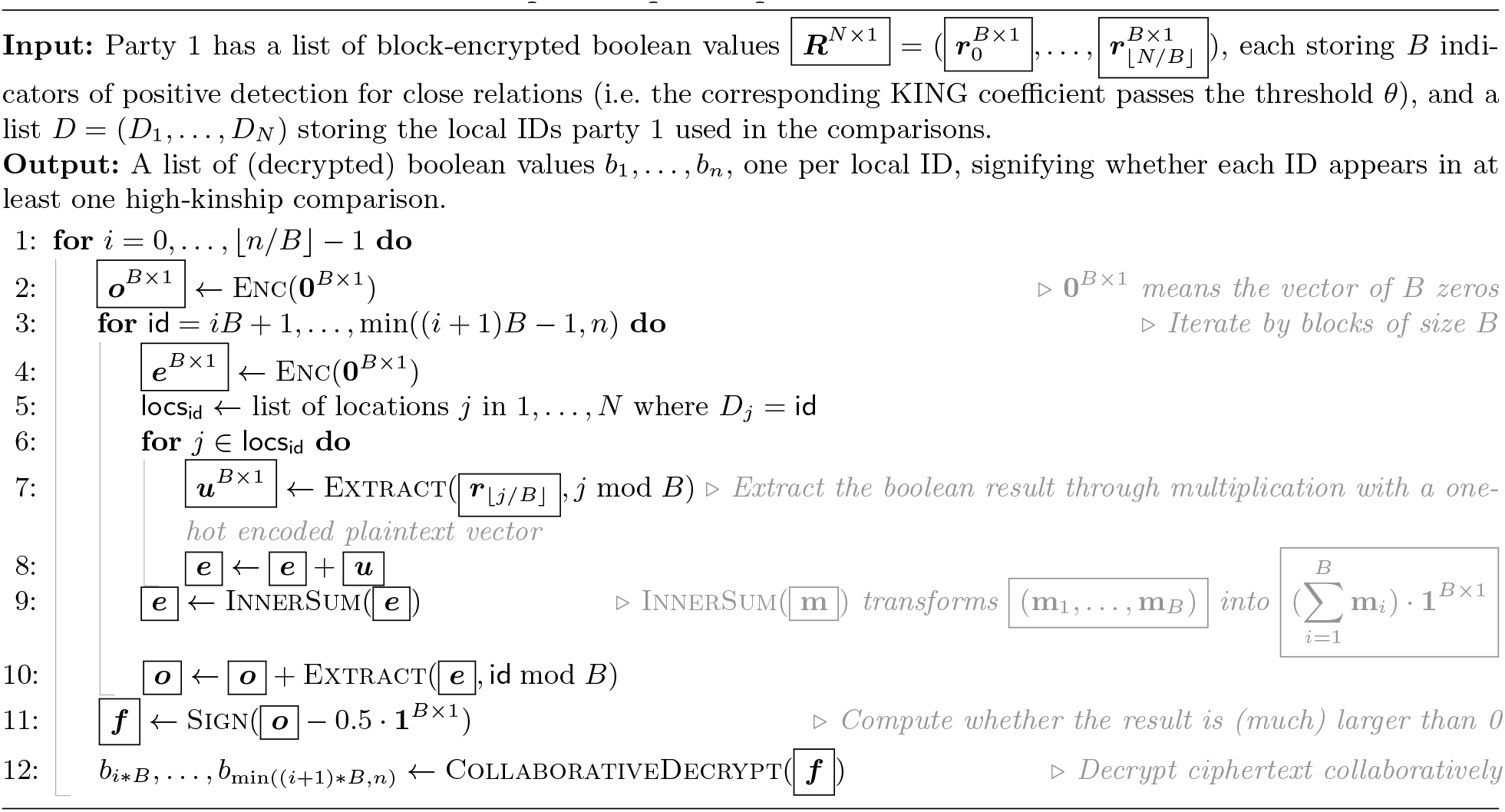

## Notes

### Competing Interest Statement

The authors have declared no competing interest.

